# Comparative analysis of genome-scale, base-resolution DNA methylation profiles across 580 animal species

**DOI:** 10.1101/2022.06.18.496602

**Authors:** Johanna Klughammer, Daria Romanovskaia, Amelie Nemc, Annika Posautz, Charlotte Seid, Linda C. Schuster, Melissa C. Keinath, Juan Sebastian Lugo Ramos, Lindsay Kosack, Annie Evankow, Dieter Prinz, Stefanie Kirchberger, Bekir Ergüner, Paul Datlinger, Nikolaus Fortelny, Christian Schmidl, Matthias Farlik, Kaja Skjærven, Andreas Bergthaler, Miriam Liedvogel, Denise Thaller, Pamela A. Burger, Marcela Hermann, Martin Distel, Daniel L. Distel, Anna Kübber-Heiss, Christoph Bock

## Abstract

Methylation of cytosines is the prototypic epigenetic modification of the DNA. It has been implicated in various regulatory mechanisms throughout the animal kingdom and particularly in vertebrates. We mapped DNA methylation in 580 animal species (535 vertebrates, 45 invertebrates), resulting in 2443 genome-scale, base-resolution DNA methylation profiles of primary tissue samples from various organs. Reference-genome independent analysis of this comprehensive dataset quantified the association of DNA methylation with the underlying genomic DNA sequence throughout vertebrate evolution. We observed a broadly conserved link with two major transitions – once in the first vertebrates and again with the emergence of reptiles. Cross-species comparisons focusing on individual organs supported a deeply conserved association of DNA methylation with tissue type, and cross-mapping analysis of DNA methylation at gene promoters revealed evolutionary changes for orthologous genes with conserved DNA methylation patterns. In summary, this study establishes a large resource of vertebrate and invertebrate DNA methylomes, it showcases the power of reference-free epigenome analysis in species for which no reference genomes are available, and it contributes an epigenetic perspective to the study of vertebrate evolution.

## Introduction

DNA methylation at the fifth carbon position of cytosines (5-methyl-cytosine) provides an epigenetic layer of genome regulation that does not involve changes in the DNA sequence. In vertebrates, DNA methylation occurs preferentially at palindromic CpG dinucleotides, where it marks both strands symmetrically. It is essential for genome integrity and contributes to the silencing of transposable elements^1^. Moreover, it is involved in the regulation of many biological processes associated with multicellular life^2^, including development^3^, differentiation^4^, and maintenance of cellular identity^5, 6^. DNA methylation has been studied extensively in the context of diseases such as cancer^7, 8^, metabolic diseases^9^, autoimmune disorders^10, 11^, and in aging^12^. From an evolutionary perspective, DNA methylation and its associated enzymes (most notably the DNA methyltransferases that “write” DNA methylation) are present throughout the animal kingdom, although it has been lost in certain species including the model organism *Caenorhabditis elegans*^13^. Supporting evolutionary roles of DNA methylation, it has been implicated in speciation^14^ and in the response to environmental influences^15, 16^.

Genome-wide DNA methylation patterns vary widely across species. Pioneering research in the early 1980s compared global levels of DNA methylation across several animal species^17, 18^, which revealed major differences between vertebrates and invertebrates^19^. Moreover, considerable variability was observed among vertebrates^20–23^. While these initial studies relied on methylation-specific restriction enzymes or on chromatography-based methods, more recent investigations used next-generation sequencing to determine DNA methylation patterns in 17 eukaryotic species (which included two vertebrates)^24^, in 13 animal species (five invertebrate and seven vertebrate species)^25^, in seven vertebrate species^26^, and in eight mammalian species^27^. High-resolution DNA methylation maps enabled initial analyses of the evolutionary relationship between DNA methylation and the underlying DNA sequence^28–30^. These previous studies were however limited to a small number of species, while an ideal study would cover many species across all branches of vertebrate evolution, such that each species becomes a complex data point in a truly integrative analysis of DNA methylation.

In human, where DNA methylation has been studied in most detail, a strong correlation exists between the genomic DNA sequence and local DNA methylation patterns^31, 32^. CpG-rich genomic regions (including many promoters and enhancers) tend to be unmethylated, except where they overlap with evolutionarily recent transposable elements^33^ or are subject to mechanisms of regulatory repression that involve DNA methylation^34^. In contrast, CpG-poor genomic regions tend to be highly methylated, except where they overlap with active transcription factor binding sites^35^ or underwent large-scale DNA methylation erosion, which is commonly observed in cancer cells and in ageing^36, 37^ The genome-wide correlation between DNA methylation and DNA sequence has enabled the prediction of locus-specific DNA methylation levels based on the underlying DNA sequence, focusing on CpG islands and gene promoters^38, 39^ and on individual CpG dinucleotides^40, 41^. Consistent with this genetic basis of DNA methylation, differences in the genomic DNA sequence between individuals have been linked to differences in DNA methylation^42^. Nevertheless, human primary samples tend to cluster according to tissue type rather than according to the sample donor^43^, indicating that DNA methylation differences between human individuals are generally less pronounced than tissue-specific differences.

To investigate DNA methylation beyond the human genome and in the broad context of vertebrate evolution, we established genome-scale DNA methylation profiles at single-base resolution across a wide range of vertebrate and invertebrate species, covering all vertebrate classes and several proximal invertebrate classes. Primary tissue samples were obtained from biobanks and other sources comprising mainly wild animals and zoo animals. We included heart and liver wherever possible, to allow for tissue-matched comparisons across species. In addition, other tissues such as lung, gills, fin, spleen, brain, lymph node, muscle, kidney, and skin were covered in a species-specific manner. Individuals were selected to prioritize healthy adults and to balance the male-to-female ratio, aiming at two to four biological replicates per species. This sampling strategy allowed us to cover a large number of species, consistent with our study’s focus on analyzing trends that hold across multiple species, rather than on the in-depth investigation of DNA methylation regulation in individual species.

DNA methylation profiling was performed using an optimized version of the reduced representation bisulfite sequencing (RRBS) assay^44–46^. Our assay enriches for CpG-rich regulatory regions but also covers many other parts of the genome, including exons, introns, intergenic regions, and repetitive elements; and it can be used to measure DNA methylation both at CpG sites and non-CpG sites in the genome. We analyzed our RRBS dataset using reference-genome independent bioinformatic methods^44^, allowing us to include many species that do not currently have a reference genome and to avoid biases due to the very different quality of available reference genomes. We previously validated this approach in a head-to-head comparison of reference-free and reference-based analysis in three species^44^. Moreover, as part of this study we confirmed by *in vitro* simulations of RRBS coverage based on existing reference genomes, comparison with whole genome bisulfite sequencing (WGBS) data, and careful analysis of potential biases, that RRBS is indeed suitable for cross-species analysis.

Our full dataset comprises 2443 DNA methylation profiles covering 580 animal species (535 vertebrates and 45 invertebrates). Based on this dataset, we identified a quantitative, predictive association of DNA methylation and the underlying genomic DNA sequence that was shared between vertebrate and invertebrate species, yet we observed two major transitions along the evolutionary axis: one between vertebrates and invertebrates and one between amphibians and reptiles. We also investigated tissue-specific and inter-individual differences in DNA methylation, and we found that tissue specificity was more pronounced in fish, birds, and mammals, while differences between individuals and between tissues had similar effects on DNA methylation variability in invertebrates, reptiles, and amphibians. By analyzing transcription factor binding sites in differentially methylated regions between heart and liver tissue throughout vertebrate evolution, we identified a deeply conserved association of DNA methylation with tissue type and cellular identity; and cross-mapping analysis identified characteristic evolutionary trends in DNA methylation at gene promoters.

In summary, this study contributes an epigenetic perspective to the investigation of vertebrate evolution, and it establishes a major resource for dissecting the role of DNA methylation in vertebrates and invertebrates. Moreover, our results underline the feasibility and value of including epigenome profiling in ongoing initiatives to map all vertebrate genomes^47^, and they provide a starting point for untangling how the complex interplay of DNA sequence patterns and DNA methylation has contributed to the evolution of vertebrate genomes.

## Results

### An atlas of DNA methylation across 580 animal species

To investigate the evolutionary dynamics of DNA methylation in vertebrates, we performed genome-scale DNA methylation profiling for 580 species and a total of 2443 primary samples (**Figure 1a-c**, **Supplementary Figure 1a-f**). Our sample collection included all vertebrate classes, and several classes of marine invertebrates, many of them closely related to vertebrates (used here as an outgroup). Specifically, we analyzed 156 samples of invertebrates (*invertebrata*), one sample of a jawless vertebrate (Japanese lamprey, *Lethenteron camtschaticum*), 32 samples of cartilaginous fish (*chondrichthyes*), 565 samples of bony fish (*actinopteri*), 74 samples of amphibians (*amphibia*), 280 samples of reptiles (*reptilia*), 607 samples of birds (*aves*), 70 samples of metatherian mammals / marsupials (*marsupialia*), and 658 samples of eutherian mammals (*mammalia*). Wherever possible, we included multiple tissues (most notably heart and liver for comparison across species) and multiple individuals, with a balanced sex ratio and a focus on young adult animals.

**Figure 1.**
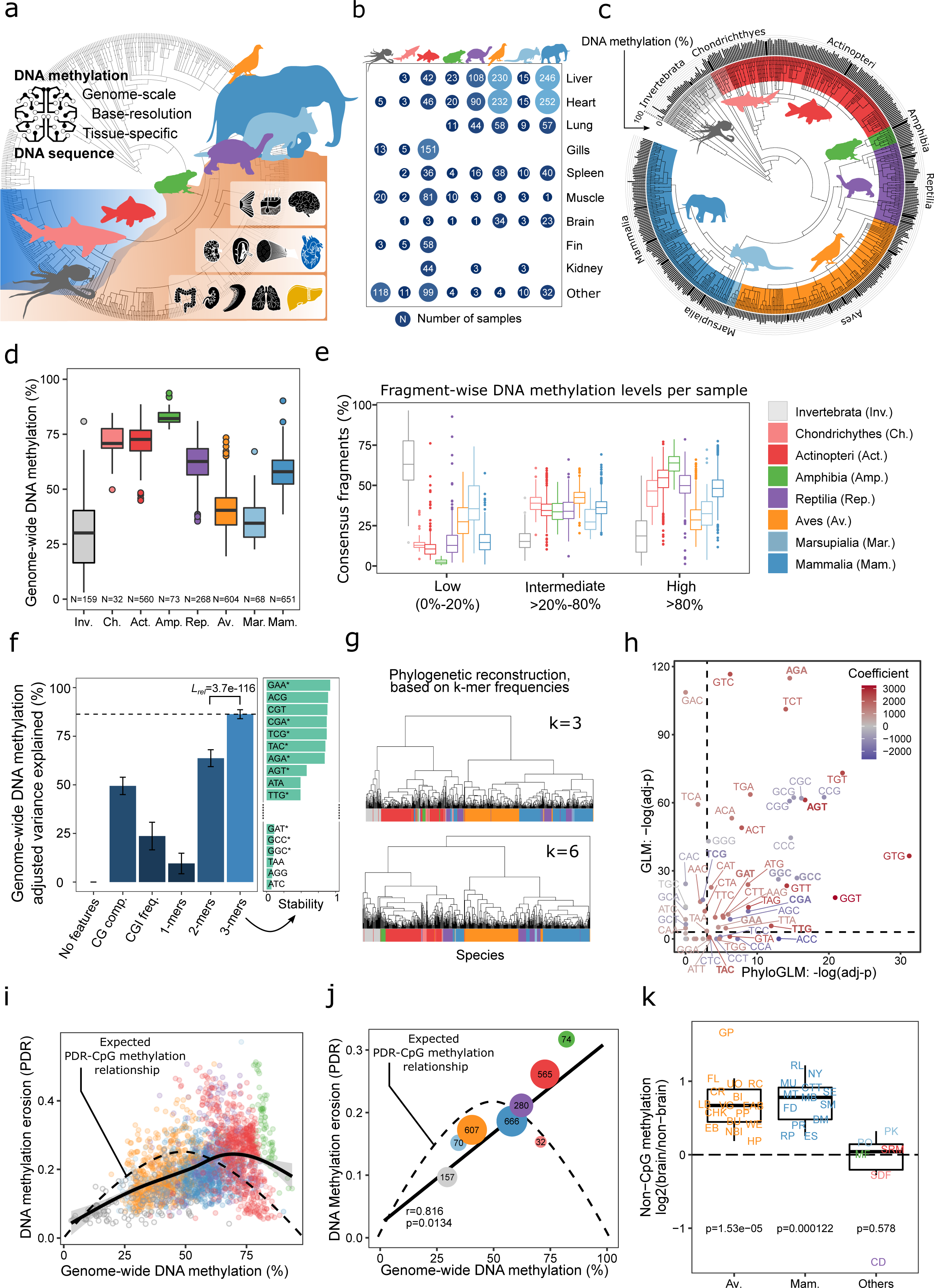
An atlas of DNA methylation comprising 583 animal species reveals global links between genomes and epigenomes throughout vertebrate evolution. (a) Visual summary of the study. The cross-species atlas comprises 2445 genome-scale DNA methylation profiles covering 583 animal species (538 vertebrates and 45 invertebrates). The animal silhouettes show one species per taxonomic group: Octopus (*invertebrata*, invertebrates), shark (*chondrichthyes*, cartilaginous fish), carp (*actinopteri*, bony fish), frog (*amphibia*, amphibians), turtle (*reptilia*, reptilians), pigeon (*aves*, birds), kangaroo (*marsupialia*, metatherian mammals / marsupials), elephant (*mammalia*, eutherian mammals). Organ silhouettes denote the main tissues included: Skin, fin, brain (ectoderm); lymph node, spleen, muscle, heart (mesoderm); gut, kidney, gills, lung, liver (endoderm). Heart and liver are highlighted in color to indicate their status as prioritized tissues in this study. Animal silhouettes were obtained from the PhyloPic database. (b) Bubble plot showing the number of analyzed primary tissues samples by taxonomic group and tissue. (c) Bar plot showing genome-wide DNA methylation levels for each species (black bars outside of the circle), averaged across all tissues and individuals, mapped onto an annotated taxonomic tree. An interactive diagram is available on the Supplementary Website (http://cross-species-methylation.computational-epigenetics.org/). (d) Boxplot showing genome-wide DNA methylation levels for all species, aggregated by taxonomic group. (e) Boxplot showing the percentage of consensus fragments in each species’ consensus reference that fall into three bins based on their DNA methylation levels, aggregated by taxonomic group. (f) Left: Bar plot showing the percentage of variance among species-specific mean DNA methylation levels that is explained by features sets reflecting genomic sequence composition (CG composition, CpG island frequency, k-mer frequencies). All values were adjusted for model complexity (i.e., number of variables), and the colors indicate the mean Akaike information criterion (AIC). Error bars represent standard deviations based on bootstrapping (100 iterations). Right: Bar plot showing the stability with which individual 3-mers were selected into the final model using stepwise selection. Stars indicate that the respective 3-mers show a statistically significant association based on the phylogenetic generalized linear model depicted in panel h. (g) Hierarchical clustering of species based on the similarity of their 3-mer and 6-mer frequencies among the consensus reference fragments. K-mer lengths of four and five yielded very similar results. The dendrogram is annotated with each species’ taxonomic group (color-coded). (h) Scatterplot comparing the statistical significance of the associations between 3-mer frequencies and global DNA methylation levels based on generalized linear models with (x-axis) and without (y-axis) correction for phylogenetic relationships. The 3-mers from panel f are shown in bold. Dashed lines correspond to an adjusted p-value of 0.05. (i) Scatterplot showing the relationship between genome-wide DNA methylation levels and DNA methylation erosion as measured by the “proportion of discordant reads” (PDR) for individual samples. The dashed line represents their mathematically expected relationship. The solid line represents a generalized additive model fitted to the data using the R function *geom_smooth*. (j) Scatterplot showing the relationship between genome-wide DNA methylation levels and DNA methylation erosion for taxonomic groups, taking the median across samples. The dashed line represents their mathematically expected relationship (as in panel i). The solid line represents a linear regression model fitted to the data. The Pearson correlation and its significance are indicated. (k) Boxplot showing log-ratios of non-CpG methylation levels in brain compared to other tissues in the same species. Boxplots are overlayed with individual data points using the species abbreviations (**Supplementary Table 2**). Increased non-CpG methylation levels in brain were assessed with a one-sided paired Wilcoxon test.

DNA methylation profiling was performed by reduced representation bisulfite sequencing (RRBS). The RRBS assay provides single-nucleotide, single-allele resolution for a defined subset of the genome, in a deterministic way that facilitates reference-free DNA methylation analysis and makes us independent of reference genomes (which are unavailable or of inconsistent quality for most analyzed species). RRBS leverages the concept of reduced representation sequencing (also known as RADseq or GBS), which is widely and successfully used for studies of evolution and ecology in species that lack a high-quality reference genome^48^. RRBS utilizes DNA methylation agnostic frequent-cutter enzymes that cut at CCGG (MspI) and TCGA (TaqI) sites, thereby enriching DNA fragments that contain at least one CpG in a species-agnostic manner and making them amenable to quantitative bisulfite-based DNA methylation analysis. RRBS covers around four out of 28 million CpGs in the human genome and two out of 20 million CpGs in the mouse genome; this includes not only CpG-rich promoter and enhancer regions, but also a broad sampling of regions with modest CpG density^49–51^.

Given extensive prior experiences with reduced representation sequencing in evolutionary studies^48^, and given the short, generic target sites of MspI and TaqI, it is unlikely that systematic differences in genome sequence composition unduly confound our RRBS profiling. To provide additional support for the validity of using RRBS for cross-species analysis, we simulated RRBS coverage across a wide range of species and analyzed the expected RRBS coverage for genomic elements such as CpG islands, transcripts, promoters, and repetitive elements. Across 76 species for which reference genomes were available (five invertebrates, one jawless vertebrate, one cartilaginous fish, eight bony fish, three amphibian, four reptiles, seven birds, three marsupial, and 44 eutherian mammals), we simulated the restriction digest and size selection in RRBS^52^, and we determined the expected coverage for each of these genomic elements (**Supplementary Figure 1b**). As expected, this analysis showed that the RRBS-specific enrichment for CpG islands was shared and consistent across all species, while systematic differences in genomic coverage between species were generally small and gradual.

Because RRBS fragments start and end at defined restriction sites, we do not depend on a reference genome or *de novo* assembly of sequencing reads; instead, we can group and overlay sequencing reads obtained from the same genomic position to construct “consensus reference fragments”. We have previously developed and extensively validated the RefFreeDMA method for RRBS-based, reference-genome independent analysis of DNA methylation^44^. Using RefFreeDMA, we combine all RRBS reads for each species into locus-specific consensus sequences with reconstructed genomic cytosines as the sites of potential DNA methylation. The resulting “consensus reference fragments” were used as the genomic reference for subsequent RRBS read alignment and DNA methylation calling, which was done separately for each sample. To be able to detect constitutively unmethylated cytosines (which appear as thymines in the RRBS reads), we also sequenced one RRBS library without bisulfite conversion for each species and included these data in the identification of genomic cytosines. RRBS quality metrics (such as the number of covered CpGs, mapping efficiency, DNA pre-fragmentation, contamination rate, conversion rate) indicated high data quality for most samples; and it allowed us to identify and flag low-quality samples (**Supplementary Figure 1g-h**; **Supplementary Table 1**).

As an additional validation, we assessed potential effects of repetitive elements, PCR amplification, and inter-individual genetic variation on our reference-free analysis (**Supplementary Figure 2a**). For each species, we empirically flagged consensus reference fragments with consistently high coverage (fourfold above average in more than 80% of samples) as likely derived from repetitive regions (“repeat”); those with sporadically high coverage (fourfold above average in less than 20% of samples) as likely subject to PCR amplification biases (“amplified”); and fragments with adequate coverage (at least half of the average) in samples from one individual but not the other individuals as likely results of genetic variation affecting the RRBS coverage (“private”). We found that the frequency of “repeat” and “amplified” fragments was generally low (below 2%) and similar across taxonomic groups, with a trend toward a lower fraction of “repeat” fragments in birds, marsupials, and mammals (**Supplementary Figure 2b-c**). We did not observe systematic effects of different PCR cycles across samples, confirming that range of cycles used (6-18) did not induce strong PCR amplification biases (**Supplementary Figure 2d**). Inter-individual genetic variation affected around 10% of the consensus reference fragments, which emphasize the importance of investigating several individuals per species.

Finally, for a subset of the species we can exploit existing reference genomes of the same or related species by cross-mapping of the consensus reference fragments (**Supplementary Figure 3a**). We pursued a data-driven approach by mapping the consensus fragments of all species to all reference genomes of animals within the same class and selected the genome with the highest mapping rate (**Supplementary Figure 3b,c**). This cross-mapping analysis was able to detect the expected association of DNA methylation with gene annotations across all taxonomic groups, including the characteristic “dip” of DNA methylation levels at the promoter region (**Supplementary Figure 3d-e**), even in extreme outliers such as the Mexican axolotl with its huge genome comprising 32 gigabases (**Supplementary Figure 3f**). We further observed that the strength of the “dip” increased with higher cross-mapping efficiency up to a rate of 25%, after which this effect leveled off (**Supplementary Figure 3g**). In aggregate, these results establish the validity of our reference-free and cross-mapping analysis of RRBS data throughout vertebrate evolution, without any concerning technical or biological biases.

**Figure 3.**
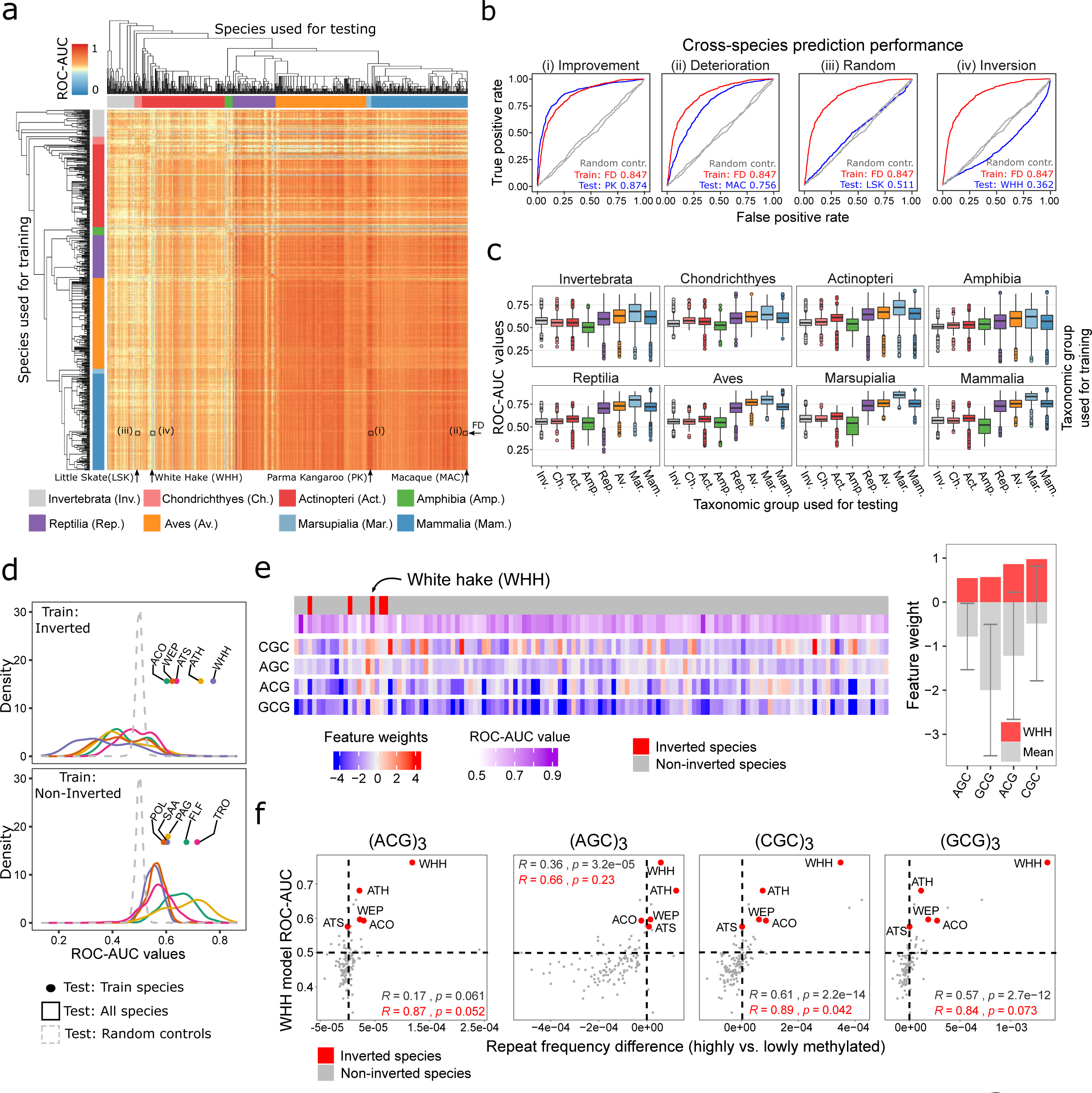
The genomic code for DNA methylation is conserved across vertebrates and invertebrates. (a) Heatmap showing ROC-AUC values for the prediction of locus-specific DNA methylation from the underlying DNA sequence between all pairs of species. SVMs were trained using data from one species (rows) and evaluated on data from another species (columns). Species were ordered by the taxonomic tree and annotated with their taxonomic group (rows and columns). The comparisons of panel b are highlighted in the heatmap. (b) ROC curves illustrating four characteristic outcomes of the cross-species predictions. SVMs were trained in one species (fat dormouse, abbreviated as FD) and tested in different targeted species (left to right: Parma wallaby, PK; macaque, MAC; little skate, LSK; white hake, WHH), resulting in: (i) high prediction performance in the target species, exceeding that observed in the training species; (ii) high prediction performance in the target species but not reaching the same level as in the training species; (iii) poor prediction performance in the target species close to the negative controls (data with randomly shuffled labels); (iv) inverted prediction performance with poorer-than-random prediction performance in the target species (“inverted species”). (c) Boxplots summarizing the cross-species prediction performance (ROC-AUC values from panel a) aggregated by taxonomic group of the training species (individual plots) and test species (x-axis). (d) Histograms of cross-species prediction performance (ROC-AUC values from panel a) for all inverted fish species (top) in comparison to the same number of phylogenetically related non-inverted species (bottom). The following inverted species are shown: Atlantic cod, ACO; walleye pollock, WEP; white hake WHH; Atlantic salmon, ATS; Atlantic herring, ATH. And the following phylogenetically related species are shown: Pollock, POL; silver arowana, SAA; Pacific grenadier, PAG; onefin flashlightfish, FLF; trout TRO. Models trained in an inverted species obtained ROC-AUC values below 0.5 in most other species, while models trained in a non-inverted species obtained ROC-AUC values above 0.5 in most other species. (e) Left: Heatmap showing SVM feature weights for the most differential 3-mers between an inverted species (white hake, WHH) and all other bony fish (*actinopteri*) species, sorted based on the taxonomic tree. Right: Bar plots for the weights of the same 3-mers in white hake compared to their average across all other bony fish (*actinopteri*) species (error bars indicate standard deviations). (f) Scatterplots for the association between the cross-species prediction performance (y-axis) of SVMs trained in an inverted species (white hake, WHH) and the difference in frequency of four 9-mer repeats (x-axis) constructed by the repetition of the differentially weighted 3-mers from panel d. Values greater than 0 indicate higher frequency in highly methylated sequences and vice versa. The following inverted species are shown: Atlantic cod, ACO; walleye pollock, WEP; white hake WHH; Atlantic salmon, ATS; Atlantic herring, ATH. Dashed lines indicate a frequency difference of 0 (vertical line) and a ROC-AUC value of 0.5 (horizontal line).

### Patterns of genome-wide DNA methylation in vertebrate evolution

Having established the validity of our dataset and analysis method, we proceeded with a systematic analysis of factors that predict genome-wide DNA methylation across the analyzed species. We calculated genomewide DNA methylation levels for each species by averaging across consensus reference fragments, tissues, and individuals, and we overlaid these species-specific aggregates with the taxonomic tree (**Figure 1c**). These values provide an assessment of DNA methylation in those areas of the genome that RRBS enriches for; they are different from the global DNA methylation levels obtained by high performance liquid chromatography measuring total 5-methyl-cytosine levels, which tend to be dominated by repetitive genomic regions^53^.

Our analysis identified lower DNA methylation levels in invertebrates compared to vertebrates, lower DNA methylation levels in birds and marsupials compared to other vertebrates, and higher DNA methylation levels in fish and amphibia compared to the other taxonomic groups (**Figure 1d**). These observations were robust across all investigated tissue types (**Supplementary Figure 4a**) and across a wide range of technical stringency thresholds (**Supplementary Figure 4b**). Moreover, we validated the identified trends on independent WGBS data for 13 species curated from the literature^25, 54–65^, with a correlation of 0.84 for genome-wide DNA methylation based on RRBS versus WGBS (**Supplementary Figure 4c**). The trends were driven by differences in the fraction of highly (greater than 80%) versus lowly (less than 20%) methylated fragments, while fragments with intermediate DNA methylation were similarly common across vertebrate classes (**Figure 1e**).

Comparing taxonomic groups, we observed strikingly lower DNA methylation for the two marsupial orders (diprotodontia and dasyuromorpha) compared to other eutherian mammals (**Supplementary Figure 4d**). The different groups of reptiles, namely lizards, snakes, turtles, and crocodiles, showed similar levels of DNA methylation – with the exception of Henophidia (a suborder of snakes including pythons and boas), which had consistently lower DNA methylation levels (**Supplementary Figure 4e**). Among the invertebrates, which are by far the most heterogeneous group in our analysis, we observed a wide range of DNA methylation levels from 2% in *Penaeus* (prawns) to 80% in *Alitta succinea* (clam worm). The majority of assessed invertebrates had DNA methylation levels between 20% and 40%, similar to those observed in birds and marsupials. Finally, *Lethenteron camtschaticum* (Japanese lamprey), which is a jawless vertebrate, showed high DNA methylation levels at 60%, which is similar to the levels observed in reptiles and mammals (**Supplementary Figure 4f-g**).

To assess the relationship between genome-wide DNA methylation levels and the underlying genome across species, we constructed linear models based on features that describe each species’ DNA sequence composition (e.g., k-mer frequencies, CG composition, CpG island frequency). Strikingly, 3-mer frequencies explained more than 80% of the observed variance in mean DNA methylation levels across vertebrate evolution (**Figure 1f**). Four of the five most predictive 3-mers contained a CpG dinucleotide (ACG, CGT, CGA, TCG), while the fifth (GAA) has been implicated in mammalian-specific repeat expansions^66^. In contrast, CpG island frequency alone explained only ∼23% of the observed variance, and CG composition (which included separate variables for C, G, and CpG frequency, as well as the CpG observed vs. expected ratio) explained around 50%.

These results show that CpG density is a key contributor but clearly not the only factor explaining the close association between genome-wide DNA methylation and genome sequence composition across species.

We found that similarities in 3-mer frequencies within the genomic DNA sequence between species closely reflect their phylogenetic distance; and the same was true k-mers of greater length as illustrated by 6-mers (**Figure 1g**). This prompted the question whether 3-mer frequencies predict genome-wide DNA methylation levels directly or through their association with phylogenetic distance. We thus compared the predictive power of 3-mer frequencies with that of phylogenetic distance, which we modeled by a representation of the taxonomic tree and by assigning each species to its corresponding taxonomic group (**Supplementary Figure 4h**). In this analysis, the prediction performance of 3-mer frequencies alone (86.4%, AIC=3819) exceeded that of both the taxonomic tree (81.0%, AIC=3622) and the taxonomic groups (74.1%, AIC=4154) alone. Nevertheless, the combination of 3-mer frequencies with phylogenetic information led to a modest further increase in overall prediction performance (87.8%, AIC=3371 for the taxonomic tree and 92.0%, AIC=3514 for taxonomic groups). We further validated these results by analyzing the links between 3-mer frequencies and genome-wide DNA methylation levels using generalized linear models that explicitly control for phylogenetic relatedness; a third of the 3-mers (22 out of 64) showed a significant association with DNA methylation beyond the effects of phylogeny (**Figure 1h**). These results support that 3-mer frequencies are directly predictive of genome-wide DNA methylation levels, beyond the strong link between phylogeny and 3-mer frequencies.

We also used our dataset to investigate DNA methylation stability and erosion in a wide range of species, motivated by recent studies that linked DNA methylation erosion to human cancers and ageing^67, 68^. We can quantify DNA methylation erosion based on our RRBS data using the “proportion of discordant reads” (PDR) metric^67^. This metric exploits that most genomic loci exhibit a bimodal distribution of DNA methylation (i.e., a locus is either fully methylated or fully unmethylated), and it interprets deviations from this pattern as evidence of DNA methylation erosion. The PDR metric was first established for cancer, where it was associated with clinical features including tumor aggressiveness^67, 69–71^. We calculated species-specific PDR values in analogy with the species-specific DNA methylation levels by averaging across consensus reference fragments, tissues, and individuals (**Supplementary Figure 5a**). We plotted the resulting values over the corresponding species’ DNA methylation level (**Figure 1i-j**), expecting high PDR values at DNA methylation levels around 50% and lower PDR values toward high and low DNA methylation levels, given the mathematical properties of the PDR metric. However, we found that DNA methylation levels of around 75% corresponded to the highest PDR values (**Figure 1i**) and that this shift was mainly driven by the taxonomic groups with high DNA methylation levels (amphibians and bony fish), as well as reptiles (**Figure 1j**). We speculate that these taxonomic groups evolved to endure higher levels of DNA methylation erosion, possibly as a consequence of high genome-wide DNA methylation levels being harder to maintain. In contrast, mammals, birds, marsupials, and cartilaginous fish showed a tendency toward lower than expected levels of DNA methylation erosion (**Figure 1j**), indicative of molecular mechanisms that foster DNA methylation stability in these groups.

We also investigated non-CpG methylation, which we can detect with RRBS as shown previously^72^. We found that non-CpG methylation was expectedly low (less than 2%) but detectable in most samples. We did not observe strong differences in non-CpG methylation across taxonomic groups – with one exception: Brain samples of birds and mammals had elevated levels of non-CpG methylation (**Supplementary Figure 5b**), which were on average 59% higher in birds and 72% higher in mammals compared to other organs (**Figure 1k**); in contrast, the difference was much weaker for the other taxonomic groups (3% higher in brain). Widespread non-CpG methylation in brain samples has been reported previously for human and mouse^73^ and has been explained by incomplete CpG specificity of mammalian DNA methyltransferases^72^. Our results suggest that this phenomenon generalizes to other mammals and birds, but is not shared by all vertebrates, which may point to differences in the DNA methylation machinery across taxonomic groups^25^.

Vertebrates have high DNA methylation levels throughout the genome, except for regions of open chromatin; in contrast, invertebrates are thought to carry a mosaic of high methylation (often at genes) and low methylation (the bulk of the genome)^74^, although some highly methylated invertebrate genomes have been described^75^. In our dataset, we observed strong differences between vertebrates and invertebrates – for genome-wide DNA methylation levels at CpGs, for DNA methylation erosion, and to a lesser degree also for non-CpG methylation (**Supplementary Figure 4a, 5a-b**). However, these differences were gradual, with overlapping distributions between vertebrate and invertebrate species, where the Japanese lamprey as a jawless vertebrate clearly sided with the vertebrates (**Supplementary Figure 5c**).

**Figure 5.**
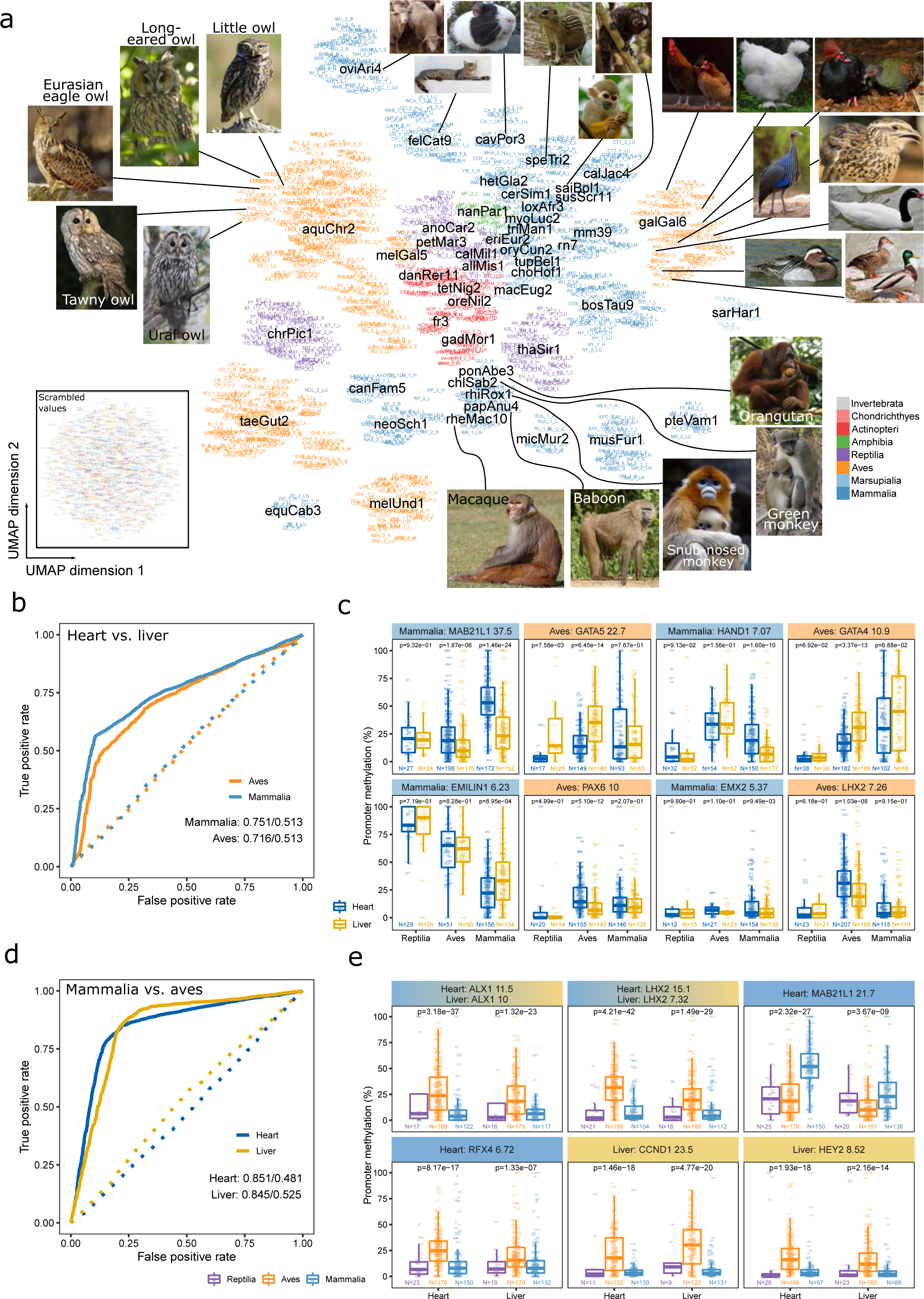
Cross-species analysis of DNA methylation in the human-ortholog gene space identifies both phylogenetic conservation and divergence of gene promoter methylation. (a) UMAP representation of DNA methylation at gene promoters based on cross-mapping of reference-free consensus reference fragments to annotated reference genomes. Samples are colored by taxonomic group, and the matched reference genomes are overlayed in black. Each sample is labeled by its sample identifier (**Supplementary Table 1**), searchable and readable when zooming into the PDF of the figure. Inset: UMAP representation of the same data but with non-missing values randomly re-assigned to non-missing positions in the data matrix to exclude potentially predictive features arising from the patterns of missing values. (b) ROC curves for random forest classifiers using the cross-mapped dataset to distinguish between heart and liver based on promoter methylation data for birds and mammals. The solid lines are based on the actual data, while the dashed lines are based on randomized data as in the inset in panel a. (c) Boxplots showing DNA methylation levels at gene promoters for the four most predictive (i.e. differential) genes in the classification of heart versus liver, aggregated by taxonomic groups and overlayed with individual data points using the species abbreviations (**Supplementary Table 2**). Gene names and corresponding classification importance are indicated. P-values were calculated using a two-sided Wilcoxon test. (d) ROC curves for random forest classifiers using the cross-mapped dataset to distinguish between birds and mammals based on promoter methylation data for heart and liver samples. The solid lines are based on the actual data, while the dashed lines are based on randomized data as in the inset in panel a. (e) Boxplots showing DNA methylation levels at gene promoters for the four most predictive (i.e. differential) genes in the classification of mammals versus birds, aggregated by taxonomic groups and overlayed with individual data points using the species abbreviations (**Supplementary Table 2**). Gene names and corresponding classification importance are indicated. P-values were calculated using a two-sided Wilcoxon test.

Finally, we investigated potential associations between DNA methylation and cancer risk, as it had been suggested previously that DNA methylation could be one of many factors that contribute to cancer suppression in large, long-lived species^76^ – for example by suppressing repetitive DNA elements that threaten genome integrity or by constraining developmental plasticity in differentiated cells. We observed a positive correlation between genome-wide DNA methylation levels and theoretical, unmitigated cancer risk of the investigated species (which we estimated based on each species’ body weight and longevity^77^). This positive association was most pronounced in birds (r=0.53) and remained statistically significant after correcting for phylogeny (p=0.01) (**Supplementary Figure 5d**). We observed similar but weaker associations also for DNA methylation erosion (likely due to positive correlation between DNA methylation and PDR) but not for non-CpG methylation. While these observations do not imply a specific causal role of DNA methylation on cancer risk, they contribute to accumulating evidence of associations between DNA methylation and cancer risk across species.

### A genomic code for DNA methylation in vertebrates and invertebrates

While the previous section focused on genome-wide measures of DNA methylation across species, our dataset also allows us to investigate locus-specific DNA methylation levels within each analyzed species, pursuing the hypothesis that there is a predictive relationship (or “genomic code”) between the DNA sequence of a given genomic region and its DNA methylation level. Previous studies in human and mouse have uncovered associations between DNA methylation and the underlying DNA sequence for genomic regions such as CpG islands and gene promoters^38, 39^ and for single CpG dinucleotides throughout the genome^40, 41^. Importantly, we use the term “code” as a shorthand for a predictive relationship between DNA sequence and DNA methylation, without implying causality or postulating a molecular mechanism that would read this code. Nevertheless, the genomic DNA sequence may encode a default epigenetic state for each genomic region, which could provide the ground state to which DNA methylation is reset in pluripotent cells and in early embryonic development^78^.

To decipher the relationship between DNA methylation and the underlying DNA sequence throughout vertebrate evolution, we trained machine learning classifiers that predict locus-specific DNA methylation levels based on the DNA sequence of the corresponding genomic regions (**Figure 2a**). Specifically, we used support vector machines with a spectrum kernel to predict the discretized DNA methylation status (highly vs. lowly methylated) of consensus reference fragments based on their genomic DNA sequence (represented by k-mer frequencies), separately for each species. The prediction performance was quantified using receiver operating characteristic (ROC) curves and area under curve (AUC) values calculated on independent test sets. The robustness of these predictions was confirmed using two alternative definitions of methylated and unmethylated regions with different stringency, resulting in highly similar ROC-AUC values (**Supplementary Figure 6a**).

**Figure 2.**
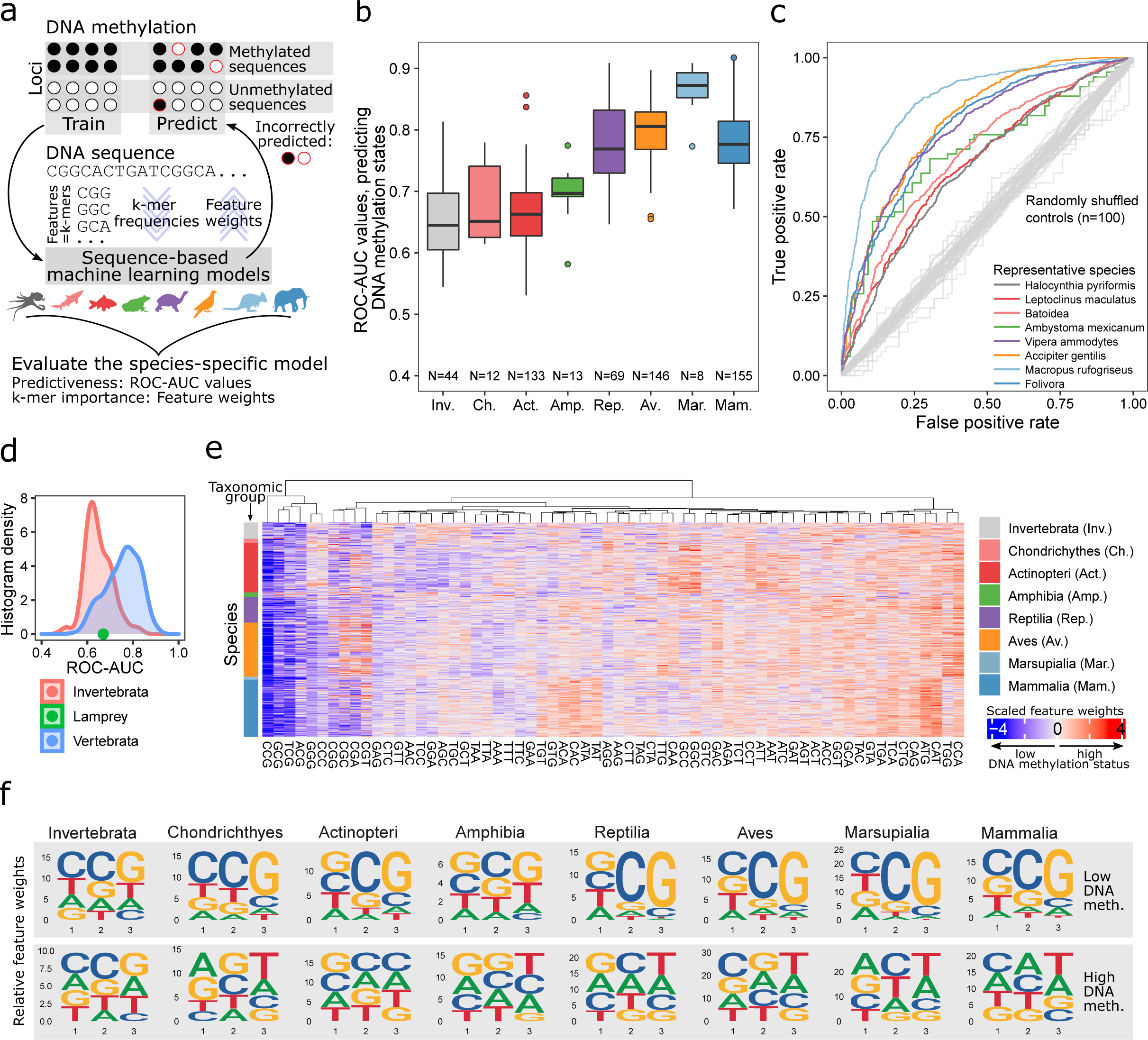
Machine learning analysis identifies a genomic code for locus-specific DNA methylation. (a) Schematic illustration of the machine learning based approach for predicting locus-specific DNA methylation from the underlying genomic DNA sequence. (b) Boxplot showing the test-set performance (operating characteristic area under curve values, ROC-AUC) of support vector machines (SVMs) predicting the DNA methylation status (high vs. low) of individual genomic regions based on the k-mer frequencies of the corresponding genomic DNA sequence. (c) Representative ROC curves for each taxonomic group, selected such that the displayed species’ ROC-AUC value closely reflects the mean ROC-AUC value of the corresponding taxonomic group. As negative controls, ROC curves trained and evaluated on data with randomly shuffled labels fall close to the diagonal (in grey). (d) Histograms of ROC-AUC values for vertebrate and invertebrate species, with the lamprey (a “primitive” jawless vertebrate) shown as a green dot between the two distributions. (e) Heatmap displaying the feature weights of 3-mers based on SVMs trained to predict locus-specific DNA methylation from the underlying DNA sequence, separately for each species (ordered by the taxonomic tree). (f) Sequence logos illustrating averaged feature weights of 3-mers across species for each taxonomic group. Sequence logos are displayed separately for positive and negative weighted features (3-mers associated with high and low DNA methylation levels, respectively).

We consistently observed better-than-random prediction performance across all taxonomic groups (**Figure 2b**, **c**), with higher ROC-AUC values in reptiles, birds, and mammals (0.78, 0.80, and 0.78) than in cartilaginous and bony fish (0.68 and 0.67) and amphibians (0.70). The prediction performance was markedly higher in marsupials (0.86) compared to eutherian mammals (0.78), indicating that DNA methylation may have a particularly pronounced genetic basis in marsupials. In contrast, the prediction performance for invertebrates (0.65) was surprisingly similar to that of fish; and the lamprey (0.67), a “primitive” jawless vertebrate, fell in between vertebrates and invertebrates (**Figure 2d**), which provides further evidence against a fundamental, qualitative difference in DNA methylation between vertebrates and invertebrates (as it exists between plants and animals)^79^. In aggregate, our results support the existence of a “genomic code” that links locus-specific DNA methylation levels to the underlying DNA sequence in vertebrate and invertebrate species.

To dissect this predictive relationship, we compared the cross-validated prediction performance of classifiers trained on 1-mer frequencies (A, C, T, G), 2-mer frequencies, etc. up to 10-mer frequencies. In this systematic analysis, 3-mer frequencies were generally the most informative, followed by 2-mer and 4-mer frequencies (**Supplementary Figure 6b-c**). In contrast, the inclusion of longer DNA sequence patterns did not result in greater predictive power, suggesting that complex sequence patterns (which may capture conserved transcription factor binding motifs) are much less relevant for the association between DNA methylation and DNA sequence than short sequence motifs. We independently validated our trained models by testing them on DNA methylation data obtained by reference-based analysis of public WGBS data for eight species. For completeness, we also performed the inverse analysis – training on WGBS data and testing in RRBS data. We observed highly consistent results, which adds further support to the validity of our RRBS-based reference-free analysis (**Supplementary Figure 6d**). We also obtained highly similar ROC-AUC values as well as preferred k-mer lengths between purely RRBS-based and purely WGBS-based predictions (**Supplementary Figure 6e**).

Finally, we inferred the predictive power of individual 3-mer frequencies for each species, and we compared the corresponding weights across all taxonomic groups (**Figure 2e-f**; **Supplementary Figure 6f**). 3-mers associated with low DNA methylation levels preferentially ended in CpG dinucleotides and started with either a C or G nucleotide. This pattern was conserved across all taxonomic groups, including invertebrates, but it was more pronounced in reptiles, birds, marsupials, and eutherian mammals compared to invertebrates, fish, and amphibians. In contrast, 3-mers associated with high DNA methylation levels followed a more balanced DNA sequence composition, and the enrichment for specific DNA sequence patterns differed between taxonomic groups. Most notably, invertebrates showed an enrichment of CpG dinucleotides also among highly methylated regions, which distinguished them from vertebrates; and mammals showed an enrichment of CpA dinucleotides, which tend to arise from the mutation of methylated CpG dinucleotides^80^.

### Conservation and divergence of the genomic code for DNA methylation

Our results support the existence of a “genomic code” or predictive relationship between DNA sequence and DNA methylation that is reflected in characteristic 3-mer frequencies of methylated and unmethylated loci and that is generally consistent across all investigated taxonomic groups. Nevertheless, we also observed characteristic differences, both between taxonomic groups and between individual species within a group. To investigate these evolutionary differences more systematically, we trained machine learning models in one species and applied them (without retraining) to predict DNA methylation levels in another species. For each pair of species, we then determined the ROC-AUC values as measures of cross-species predictability, with high values indicating good transferability of the trained model between the pair of species (**Figure 3a-b**).

The prediction performance was generally high between related species and even across taxonomy groups (**Figure 3c**), often reaching similarly high values as for the prediction within a species or within a taxonomy group. However, we also observed pronounced differences in cross-species predictability, notably between invertebrates, fish, and amphibians on the one hand (where the predictability was lower) and the evolutionarily younger groups of reptiles, birds, and mammals on the other hand (where the predictability was higher).

We found that models trained in species with lower prediction performance generally performed well in species with higher prediction performance, but not vice versa (**Figure 3a-c**). Even prediction models trained in invertebrates retained some predictive power in vertebrates, despite fundamental differences in the genomic distribution of DNA methylation between vertebrate and invertebrate genomes^81^. These observations suggest that the predictive relationship between locus-specific 3-mer frequencies and their associated DNA methylation levels (i.e., the “genomic code” of DNA methylation) is deeply conserved across all taxonomic groups. However, the fact that predictability of DNA methylation differed widely across target species suggests that some species deviate much more strongly from the genetically encoded “default” DNA methylation profile than other species (e.g., due to tissue-specific regulation, environmental influences, and/or stochastic effects).

Curiously, a few species of invertebrates, fish, amphibians, and reptiles showed an apparent inversion of the genomic code for DNA methylation, such that DNA sequence patterns normally associated with low DNA methylation levels were instead linked to high DNA methylation levels and vice versa. The existence of such inverted species in our dataset was evident from cross-species ROC-AUC values in the target species that were substantially worse than expected by random chance (blue horizontal/vertical stripes in **Figure 3b**). In other words, prediction models trained in non-inverted species and applied in inverted species misclassified methylated regions as unmethylated, and unmethylated regions as methylated, with frequencies that could not be explained by random chance. We observed the same pattern of significantly worse-than-random prediction performance when models were trained in inverted species and applied in non-inverted species. In contrast, cross-species prediction between two different inverted species were generally more coherent with each other than with non-inverted species (**Supplementary Figure 7a**), suggesting a shared underlying mechanism.

To investigate the biological basis of the apparent inversion of the genomic code for DNA methylation, we focused on the white hake (*Urophycis tenuis*), which is one of the bony fish (*actinopteri*) species with a pronounced inversion (**Figure 3d**) that was consistently detected across tissues and individuals (**Supplementary Figure 7b-f**). Comparing the feature weights of the machine learning classifiers, we identified four 3-mers (AGC, GCG, ACG, CGC) that were strongly predictive of high DNA methylation levels in the white hake sample but predictive of low DNA methylation levels in other bony fish (**Figure 3e**). These 3-mers were associated with repetitive elements in the white hake and to a lesser degree also in other inverted fish species, but not in most of the non-inverted fish species (**Figure 3f**). We thus conclude that the observed inversion may be explained by introgression of evolutionarily recent, CpG-rich, repetitive elements, which tend to acquire high DNA methylation levels as part of the cells’ machinery for suppressing their instability and expansion^1^.

### Conservation of DNA methylation patterns underlying tissue type

The hypermethylation of repetitive elements in the inverted species (as described in the previous section) appears to occur on top of a broadly conserved “genomic code” of DNA methylation; and we would expect that the same applies to tissue-specific as well as inter-individual differences in DNA methylation. To investigate the relative contributions of the tissue and the individual to the DNA methylation variation in our dataset, we focused on those species for which we have multiple tissues and individuals (n=360), and for each species we inferred the percentage of variance explained by the tissue and by the individual (**Figure 4a**).

**Figure 4.**
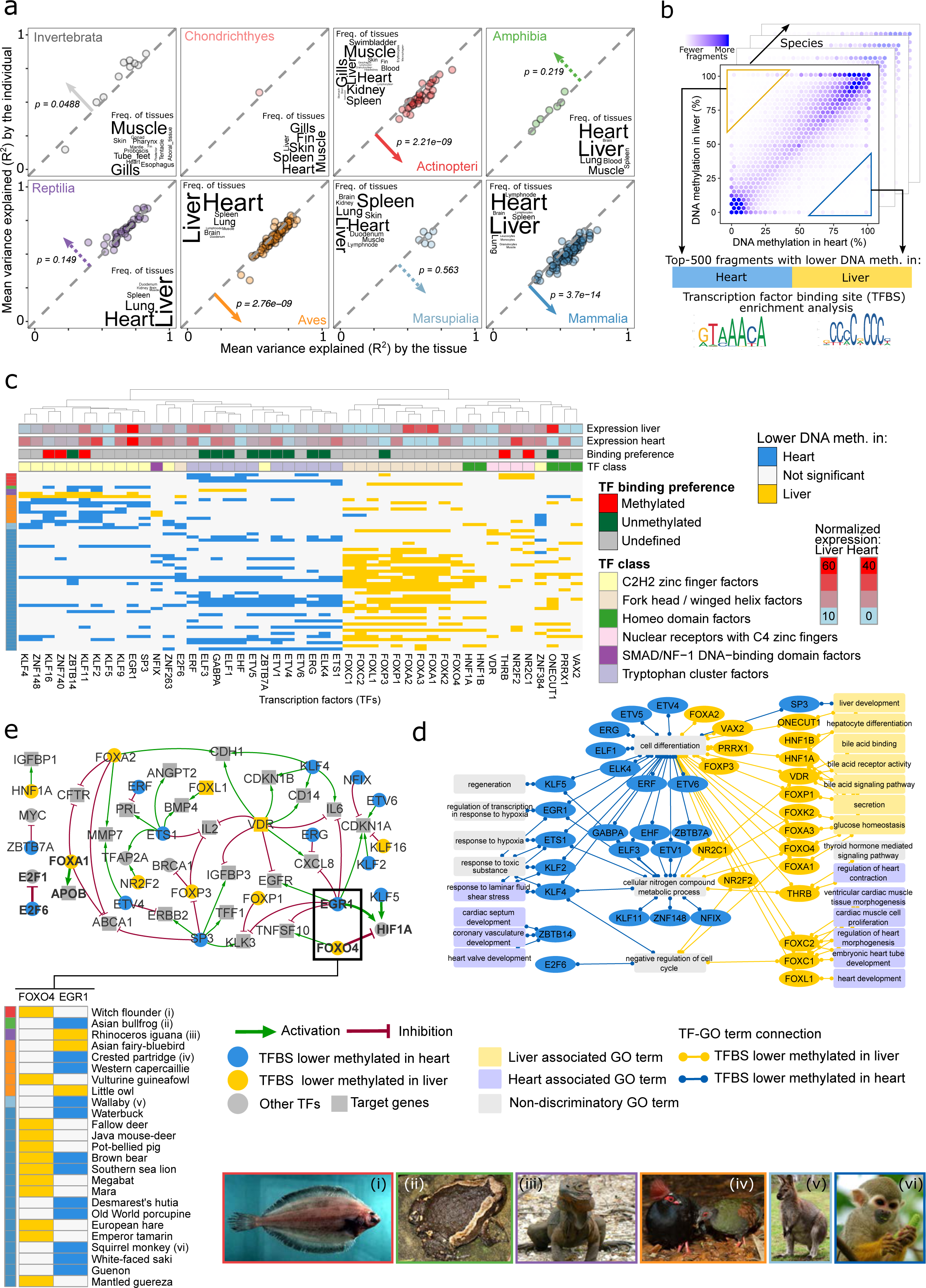
Tissue-specific DNA methylation analysis indicates deeply conserved associations of DNA methylation with transcription regulation and cellular identity. (a) Scatterplots showing the percentage of locus-specific DNA methylation variance that is explained by the tissue (x-axis) and by the individual (y-axis), separately for each taxonomic group. Arrows and p-values indicate the direction and statistical significance of the difference in the variance explained by tissue and individual, calculated using a two-sided pairwise Wilcoxon test. Dashed lines indicate non-significant differences. Word clouds summarize the frequency of tissue types that contributed to the analysis in each taxonomic group. (b) Schematic illustration of the enrichment analysis for transcription factor binding site (TFBS) motifs among the differentially methylated regions identified between heart and liver (within a given species). (c) Clustered heatmap showing TFBS motif enrichments for differentially methylated fragments between heart and liver. For each transcription factor (columns), colors indicate whether it was enriched in fragments that were hypomethylated in heart (blue) or liver (yellow) in the corresponding species (rows). Only transcription factors and species with a minimum of ten significant enrichments per species and normalized RNA expression values greater than one in either heart or liver tissues in the Human Protein Atlas^96^ are shown. (d) Visualization of the Gene Ontology annotations of the transcription factors identified in panel c. (e) Gene-regulatory network constructed based on the transcription factors identified in panel c with known binding preference (methylated/unmethylated) and their direct target genes with known regulatory effect (activation: green; repression: red). Transcription factors that were preferentially hypomethylated in one tissue type were colored in yellow (heart) or blue (liver), while those that did not show such an enrichment as well as the transcription factor target genes were colored in grey. The inset shows the specific enrichments for FOXO4 and EGR1 in heart and liver, which have opposing effects on HIF1A (FOXO4: activation; EGR1: repression). The pictures at the bottom show one species for each taxonomic group that contributed to this cross-species analysis of DNA methylation differences in heart and liver.

In human and mouse, it is well established that DNA methylation patterns are more similar among samples of the same tissue from different individuals than among samples of different tissue from the same individual^43, 82, 83^. We observed this pattern for selected species across all taxonomic groups except cartilaginous fish (**Supplementary Figure 8a**). However, when quantifying this phenomenon across all species, we found that tissue-specific differences clearly exceeded inter-individual differences only in mammals, birds, and bony fish, whereas we observed equal or higher variability explained by individual than by tissue for many invertebrates, amphibians, and reptiles (**Figure 4a**). This observation was not due to differences in technical data quality, as measured by PCR enrichment cycles in the RRBS protocol (**Supplementary Figure 8b**) and DNA pre-fragmentation as a proxy for low DNA quality (**Supplementary Figure 8d**), although we observed a slight negative association between DNA pre-fragmentation and the overall variance explained. We also investigated potential confounding effects of genetic variation within each species, which we estimated based on the mean overlap in covered CpGs between samples from the same species. We did not observe particularly low concordance levels in invertebrates, amphibs, or reptiles (**Supplementary Figure 8c**), arguing against genetic variation as a major confounder in our analysis. Finally, we observed a negative correlation between DNA methylation erosion and total variance explained in reptiles, birds, and mammals (**Supplementary Figure 8e**).

Next, we sought to establish an initial concept of DNA methylation in its relation to tissue identity – not in one specific species but across vertebrate evolution, with each species contributing a data point. We focused on heart and liver, including 207 species with samples from at least two individuals for both tissues. For each species, we identified differentially methylated consensus reference fragments between heart and liver (**Figure 4b**) and compared the enrichment for transcription factor binding motifs between fragments with lower DNA methylation levels in heart vs. liver, and vice versa (**Figure 4c**, **Supplementary Figure 9a**). This analysis exploits that many transcription factors and their binding motifs are conserved across vast evolutionary distances^84^. We indeed detected many transcription factor binding motifs at similar frequencies in fragments from all taxonomic groups, with no obvious preference for mammals (**Supplementary Figure 9b**).

For further biological interpretation, we determined the transcription factors that are expressed in human heart or liver tissues and whose binding sites were enriched in differentially methylated fragments, and we annotated them with GO terms related to heart and liver biology, physiology, and gene regulation (**Figure 4d**). We found that transcription factors associated with fragments characterized by lower DNA methylation levels in heart were preferentially annotated with heart-specific biological functions (e.g., ZBTB14 has a role in cardiac septum development; KLF4, KLF2 as well as ETS1 are involved in the response to laminar fluid shear stress). Conversely, fragments with lower DNA methylation levels in liver were annotated with liver-specific functions (e.g., ONECUT1 and HNF1A contribute to liver development; FOXP1, FOXA1, FOXK2, FOXA3, and FOXO4 are involved in glucose homeostasis). Moreover, the binding sites of several transcription factors with a role in the response to hypoxia and to toxic substances had lower DNA methylation levels in heart than in liver, which may be linked to the liver’s greater tolerance to such exposures. While these results are consistent with the traditional concept that low DNA methylation levels are associated with high regulatory activity^35^, we also found one striking example in which higher DNA methylation levels appear to coincide with higher regulatory activity: Fragments enriched for the binding sites of FOXC2, FOXC1, and FOXL1 – three FOX family transcription factors with an established role in heart development – showed lower DNA methylation levels in liver than in heart, indicating more diverse relationships between DNA methylation and regulatory activity^35^.

Finally, we inferred the evolutionary conserved “tissue of activity” for individual transcription factors by transcription factor binding site enrichment, while taking into account preferential binding to unmethylated or methylated DNA^85^ (**Figure 4c**). From the identified transcription factors we derived a gene-regulatory network using regulator interactions obtained from the TRRUST v2 database^86^ (**Figure 4e**). This network constitutes a first exploratory attempt at reconstructing a deeply conserved basis of the epigenetic cell identity for heart and liver tissue across vertebrates. This analysis suggests that FOXA1 (also known as hepatocyte nuclear factor 3-alpha) is active in liver, possibly inducing APOB, a crucial component of low density lipoprotein (LDL) produced in the liver and small intestine^87^. E2F6, a repressive transcription factor involved in cell cycle regulation^88^, showed higher inferred activity in heart compared to liver, potentially reflecting the very different regenerative potential of these two organs. HIF1A (Hypoxia-inducible factor 1-alpha) may be repressed by high activity of FOXO4 in liver, while being activated by KLF5 and EGR1 in heart, which might contribute to the higher tolerance toward hypoxic conditions in the liver^89^. These observations were in line with our GO analysis (**Figure 4d**) and suggest that DNA methylation may help stabilize the fundamental regulatory processes underlying vertebrate tissue identity in ways that are conserved across large evolutionary distances.

### Gene-centric patterns of DNA methylation in vertebrate evolution

The reference-free DNA methylation analysis of DNA methylation (as described in the previous sections) allowed us to include all 580 species, unconstrained by the availability of reference genomes. However, this approach makes it difficult to link DNA methylation to the genes and promoters that it may regulate. We therefore complemented our reference-free analysis of tissue-specific DNA methylation with a reference-based analysis in which we cross-mapped the samples to annotated reference genomes of the same or related species, with data-driven selection of the most fitting reference genome (**Supplementary Figure 3, 10a**). We calculated mean DNA methylation levels of individual gene promoters for each sample, based on the gene annotations of the corresponding reference genome. We then linked these annotations to their human ortholog, to analyze gene promoter methylation from all species in one shared gene space (**Supplementary Figure 10a**). We thus derived a gene-centric DNA methylation landscape comprising 382 species, 1524 cross-mapped samples, and 14,339 genes. We projected the samples into two dimensions using the UMAP method (**Figure 5a**).

This cross-species landscape of DNA methylation at gene promoters reflects phylogenetic relationships at the level of taxonomic groups and reference genomes, together with more fine-grained patterns determined by species, tissues, and individuals (**Figure 5a**, **Supplementary Figure 10b**). The highest resolution was obtained for mammals, given the large number of available reference genomes and high conservation of human genes ensuring an accurate mapping. Samples mapped to reference genomes of old world monkeys (rhesus, baboon, snub-nosed monkey, green monkey) and apes (orangutang) formed a concise cluster, while samples mapped to the reference genomes of new-world monkeys (marmoset, squirrel monkey) formed a separate group, supporting that our method captures genetic and epigenetic similarity without undue bias due to certain reference genomes. Among the birds, the golden eagle genome (genome assembly: aquChr2) and the chicken genome (genome assembly: galGal6) enabled gene-centric analyses for multiple other species including owls and ducks, respectively. Fish, amphibians, and reptiles were not as well represented as mammals and birds but still detectable in this gene-centric analysis, exploiting high conservation of certain genes across long evolutionary timescales. The observed patterns were clearly non-random and not seen in scrambled data (**Figure 5a, inset**).

We exploited this cross-species landscape to define groups of genes that exhibit similar patterns of DNA methylation at their promoters throughout vertebrate evolution. To this end, we projected all adequately covered genes into two dimensions using the UMAP method, and we identified five distinct gene sets using the Leiden clustering method (**Supplementary Figure 10c**). Cluster 1 was characterized by high promoter methylation in mammals, and specifically in samples from the lymph node (a tissue that is largely restricted to mammals); Cluster 2 showed consistently low promoter methylation across taxonomic groups and tissues, and was enriched for GO terms related to organ morphogenesis; Cluster 3 exhibited high levels of promoter methylation in birds, and in brain and several internal organs, and it was enriched for GO terms relating to organism development; Cluster 4 was associated with high promoter methylation in reptiles and bony fish but low promoter methylation in cartilaginous fish; Cluster 5 was characterized by low promoter methylation in various internal organs but high promoter methylation in blood, skin, fins, and gonads (**Supplementary Figure 10d-e**).

To assess evolutionary conservation as well as divergence of tissue-specific DNA methylation for individual gene promoters, we employed random forest classification for robust identification of differentially methylated genes across species (**Figure 5b-e**). We focused on the two best represented tissues (heart, liver) and the two best represented taxonomic groups (mammals, birds) and devised four classification tasks: Heart versus liver tissues in each of the two taxonomic groups, and mammals versus birds for each of the two tissues. In these analyses, we ensured that all models were tested only on species that had not been used during training, in order to focus these analyses on patterns that are conserved across species. We found good prediction performance for all four tasks: ROC-AUC in the heart versus liver classification were 0.751 for mammals and 0.716 for birds, while the corresponding values for the classification of mammals versus birds were 0.851 for heart and 0.845 for liver (**Figure 5b-c**).

We investigated which gene promoters enable these predications and found that the most discriminatory genes between heart and liver (**Figure 5c**) were transcription factors. In birds, this included GATA4 and GATA5, both with well-known roles in heart differentiation. In mammals, in addition to three transcription factors (MAB21L1, HAND1, EMX2), we identified the EMILIN1 gene, which codes for a protein that anchors smooth muscle cells to elastic fibers potentially relevant for heart function. This gene showed increasing promoter methylation from mammals to reptiles, possibly relating to the marked anatomical and function changes during the evolution of vertebrate circulatory systems. Moreover, MAB21L1, a cell fate regulator with similarity to the cGAS innate immune sensor^90^ showed higher promoter methylation in heart tissues of both mammals and birds. The most discriminatory genes between birds and mammals (**Figure 5e**) were similar between heart and liver tissue, for example the homeobox genes ALX1 and LHX2 or the cell cycle promoting cyclin CCND1, which showed significantly higher promoter methylation levels in birds compared to reptiles and mammals.

Finally, we performed gene-centric analyses of promoter methylation across all eight taxonomic groups (**Supplementary Figure 10f**), and we identified 48 genes that had a conserved promoter methylation signal across most of the assessed taxonomic groups. Only one gene retained an unmethylated promoter throughout vertebrate evolution: CHCHD7, a putative housekeeping gene that is ubiquitously expressed in human tissues. In contrast, the promoter of SPON2, which codes for a cell adhesion protein involved in innate immunity, was highly methylated in all classes expect marsupials. Genes with high promoter methylation across taxonomic groups (such as SPON2, LMF1, NRDE2, SLC38A10, VASN, NUDT7, GNL2, NETO1, APRT, FAM163B, ALOX5) had a tendency toward higher methylation levels in mammals, while most other genes had low DNA methylation levels in mammals. A similar pattern was observed also for reptiles, while birds and marsupials often showed low promoter methylation values even for genes with highly methylated promoters in other taxonomic groups. Fish and amphibia showed high promoter methylation in most of the 48 broadly conserved genes, consistent with their generally high DNA methylation levels. Lamprey, as a jawless vertebrate, showed promoter methylation patterns similar to those of cartilaginous fish. In contrast, invertebrates generally showed low promoter methylation levels even for genes with high promoter methylation across all other taxonomic groups, supporting diverging gene-regulatory roles of DNA methylation between vertebrates and invertebrates.

Given the breadth of the presented dataset and analysis, detailed follow-up studies in selected species will be needed to corroborate and extend these observations; and we provide our dataset as a comprehensive resource and starting point for such investigations (http://cross-species-methylation.computational-epigenetics.org/).

## Discussion

DNA methylation has important roles for genome integrity, regulation of gene expression, and cellular identity. Previous reports indicated that the genomic distribution and biological functions of DNA methylation systematically differ between vertebrate and invertebrate model organisms (a view that has recently been challenged^75, 91^). However, a systematic analysis of DNA methylation patterns throughout vertebrate evolution has been lacking. We thus established a large dataset of genome-scale, single-nucleotide DNA methylation profiles for 2445 primary tissue samples covering 580 animal species (535 vertebrates and 45 invertebrates). The size and resolution of this dataset allowed us to address fundamental biological questions with adequate resolution and statistical power, including the predictive relationship of DNA methylation and DNA sequence, prevalence of DNA methylation erosion, role of tissue versus individual as sources of DNA methylation variation, and conservation of gene-regulatory DNA methylation signatures throughout vertebrate evolution.

This study was enabled by a highly scalable method for DNA methylation profiling and data analysis that is applicable to essentially any species and tissue, allowing us to capitalize on large zoological biobanks and sample collections of wild, pet, and zoo animals. The RRBS assay proved robust across variable DNA quantities and qualities, consistent with our previous experience with challenging formalin-fixed paraffin-embedded patient samples^69^; this facilitated the inclusion of primary tissue samples obtained from deceased animals (65% of analyzed samples were collected as part of routine animal pathology). Using our RefFreeDMA software, we were able to analyze and compare DNA methylation between tissues, individuals, and species, independent of whether a reference genome has been established for these species. We extensively validated the RRBS assay and the reference-free analysis pipeline with coverage simulations across 76 reference genomes and validation of key results by a meta-analysis of WGBS data for 13 species. In addition, we cross-mapped the results of our reference-free analysis to gene-annotated reference genomes, which provided additional validation and gene-centric insights, while also illustrating the future value of our dataset for reference-based analysis as many more high-quality reference genomes will become available over the next decade.

At the core of our study, we used machine learning to associate DNA methylation with the underlying DNA sequence, thereby linking aspects of genomes and epigenomes throughout vertebrate evolution. We refer to the predictive relationship between DNA methylation and the underlying DNA sequence as a “genomic code” that links methylated as well as unmethylated states to preferred sequence motifs, without implying any specific mechanism or direct causation. We found that this “genomic code” was highly conserved across all analyzed taxonomic groups. Both for genome-wide and locus-specific DNA methylation levels, this relationship was best described by 3-mer frequencies. As expected, high frequency of CpG dinucleotides was associated with low DNA methylation levels, but CpGs were by no means the only contributing factor. Machine learning models trained to predict the locus-specific DNA methylation level from the underlying DNA sequence in one species generally performed well also in other species, even across taxonomic groups, and the prediction performance appeared to be more a feature of the target species than of the species in which the model was trained.

Surprisingly, the “genomic code” was detectable even among invertebrate species, to the point that models trained on DNA methylation data for invertebrate species retained some predictive power in vertebrate species. More generally, our dataset uncovered an unexpected degree of conservation in the characteristics of DNA methylation between vertebrates and invertebrates. First, while invertebrates on average showed lower genome-wide DNA methylation levels than vertebrates, many invertebrate species exhibited genome-wide DNA methylation levels well within the distribution of vertebrates (**Supplementary Figure 4f**, **5c**). Second, certain invertebrate species including sea urchins (Strongylocentrotus) showed the typical DNA methylation profiles of vertebrates, with a prominent dip at gene promoters and high gene-body methylation (**Supplementary Figure 3d-e**). Third, the lamprey fell between vertebrates and invertebrates in terms of the predictiveness of the “genomic code” of DNA methylation, consistent with its intermediate position as a jawless vertebrate.

These results support that the changing characteristics of DNA methylation throughout vertebrate evolution are more gradual and diverse than they appeared based on previous analyses of much fewer model organisms. Nevertheless, two major transitions in the “genomic code” of DNA methylation are supported by our dataset: With the emergence of vertebrates and the emergence of reptiles. These transitions manifested themselves not only through increased predictability of DNA methylation from DNA sequence (prediction accuracies were generally higher for reptiles, birds, marsupials, and mammals than for fish and amphibians), but also in the similarity of predictive sequences across species within taxonomic groups and in a shift toward higher predictive power of CpG-rich 3-mers for lowly methylated loci (**Figure 2**). We speculate that this “genomic code” may play a role in restoring default DNA methylation patterns not only in embryogenesis^78^, but also following artificial DNA methylation depletion^92, 93^. This DNA sequence encoded default epigenetic state could provide a basis that is modulated by other effects (e.g., tissue, environment, and random chance) over the course of an animal’s development and life. While this study was not designed to elucidate a potential mechanistic basis of the described “genomic code”, it will be interesting to combine our cross-species data with an investigation of the biochemical machinery that controls DNA methylation – including DNA methyltransferases^13^ and demethylases^94^, but also histone-modifying enzymes^95^ and transcription factors that modulate DNA methylation.

A side note of our study is the cross-species analysis of DNA methylation erosion, which uncovered high levels of DNA methylation erosion in the taxonomic groups with the highest genome-wide DNA methylation levels (amphibians and fish), rather than in the species with intermediate DNA methylation levels (mammals and reptiles) as we would mathematically expect^67^. This observation may be due to higher DNA methylation levels being intrinsically harder to maintain, given the limited fidelity of maintenance DNA methylation^96^. The affected species would therefore be able to tolerate lower stability of DNA methylation, with the potential upside of creating more room for accommodating environmental influences on DNA methylation.

Erosion of DNA methylation patterns has been observed in human cancers^67, 69–71^, and loss of epigenetic control appears to causally contribute to cancer development^8, 97, 98^. In this context, the observed variability in DNA methylation erosion between taxonomic groups raises interesting questions regarding a potential role of DNA methylation in the protection against cancer risk, especially in large and long-lived vertebrates. While our dataset cannot conclusively address such questions, we observed intriguing associations such as low levels of DNA methylation erosion and a negative correlation between theoretical cancer risk and DNA methylation levels in birds, which have a low incidence of tumors^99^. We envision that our optimized RRBS assay and reference-free analysis will facilitate DNA methylation profiling of tumors of wild and captive animals for a wide range of vertebrate species encountered in veterinary pathology. This will in time contribute to a better understanding of the potential roles of DNA methylation in solving the lack of correlation between body size and cancer risk (Peto’s paradox)^100, 101^, which stands out as a remarkable feat of vertebrate evolution.

Potential limitations of this study arise from the experimental choices that allowed us to process 2443 primary tissue samples from 580 species. First, RRBS uses restriction enzymes to pre-enrich a “reduced representation” of the genome prior to bisulfite conversion and sequencing. Compared to WGBS, RRBS covers fewer CpGs (mean: 2.5 million CpGs per sample), is cheaper and more scalable; and it provides consistent starting points for the DNA fragments (i.e., the restriction sites), which facilitates the comparison between tissues and individuals (mean: 1.7 million shared CpGs across samples). Second, analyzing any subset of CpGs in the genome bears the risk of introducing species-specific biases; while we perform extensive validations and designed our analyses to ameliorate this risk, it is a relevant consideration for all analyses of the presented dataset. Third, we focused our initial analysis of this large dataset primarily on DNA methylation at CpG dinucleotides, given its well-established biological roles. Nevertheless, the RRBS assay also covers DNA methylation at non-CpG sites (i.e., CpA, CpC, CpT), and we observed expectedly low levels of non-CpG methylation in our dataset (species medians ranging from 0.99% to 2.43% across all analyzed vertebrate species). We also detected significantly higher non-CpG methylation levels in brain compared to other tissues in mammals as well as birds, consistent with a recent report focusing on much fewer species^25^. Fourth, this study relies on our reference-free analysis method (RefFreeDMA)^44^, which enables us to work without reference genomes but lacks the regional context that is provided by a high-quality reference genome. We addressed this limitation by focusing on transcription factor binding sites, whose DNA methylation levels tend to reflect the activity of the corresponding transcription factors. Moreover, we devised a cross-mapping strategy that leverages gene annotations from existing reference genomes and combines different species in a human ortholog gene space. Fifth, despite the large number of samples and species in our study, many interesting clades (especially among amphibians and reptiles) remain underrepresented or are covered with few samples. Finally, the different species do not provide fully independent data points but are connected through evolution, which we accounted for by statistical methods that correct for phylogenetic relationships or within-species modeling. Accounting for phylogenetic relationships will be an important consideration for future analyses building on our dataset, and the development of phylogenetically aware machine learning methods could refine and enhance the inferred “code”.

In conclusion, this study provides an initial account of the DNA methylation dynamics associated with vertebrate evolution, not only by creating a dataset of unprecedented scale and scope, but also by providing insights into the conserved and divergent role of DNA sequence composition, tissue types, and inter-individual variation on DNA methylation. Most notably, we found that DNA sequence and DNA methylation maintained complex associations throughout vertebrate evolution, which likely contribute to the diversity of epigenetic regulatory processes observed in vertebrate species, human populations, and complex diseases.

## Acknowledgements

We thank the Biomedical Sequencing Facility at CeMM for assistance with sequencing and all members of the Bock lab as well as Matthew Huska for their help and advice. J.K. was supported by a DOC Fellowship of the Austrian Academy of Sciences. C.B. was supported by a New Frontiers Group award of the Austrian Academy of Sciences as well as a European Research Council (ERC) Starting Grant (no. 679146) and Consolidator Grant (no. 101001971) of the European Union’s Horizon 2020 Research and Innovation Programme. The computational results have been obtained in part using the Vienna Scientific Cluster (VSC).

Animal images provided in the paper are either personal contributions from the authors or obtained from the Wikimedia Commons and other sources listed below. We thank the photographers for providing their photos under the Creative commons license CC BY-SA 3.0 (https://creativecommons.org/licenses/by-sa/3.0/deed.en):

Figure 4:

Witch Flounder: NOAA Fisheries (https://www.fisheries.noaa.gov/species/witch-flounder)

Asian banded bullfrog: D. Gordon E. Robertson (https://animalia-bio.us-east-1.linodeobjects.com/animals/photos/full/original/banded-bullfrog-28kaloula-pulchra292c-angkor-wat2c-cambobia.webp)

Rhinoceros Iguana: Tim Ross (https://commons.wikimedia.org/wiki/File:RhinoIguanaMay07Pedernales.jpg)

Crested Wood Partridge: Brian Gratwicke (https://flickr.com/photos/19731486@N07/5110329309)

Red necked wallaby: benjamint444 (https://commons.wikimedia.org/wiki/File:Red_necked_wallaby444.jpg)

Squirrel Monkey: Luc Viatour (https://commons.wikimedia.org/wiki/File:Saimiri_sciureus-1_Luc_Viatour.jpg)

Figure 5:

Sheep: Keith Weller (http://www.ars.usda.gov/is/graphics/photos/apr12/k4166-5.htm)

Guinea pig: Plath81 (https://en.wikipedia.org/wiki/Guinea_pig#/media/File:George_the_amazing_guinea_pig.jpg)

Cat: Alvesgaspar (https://commons.wikimedia.org/wiki/File:Cat_August_2010-4.jpg)

Thirteen-lined ground squirrel: Phil Myers, Museum of Zoology, University of Michigan (http://www.genome.gov/pressDisplay.cfm?photoID=4)

Squirrel monkey: Luc Viatour (https://en.wikipedia.org/wiki/Squirrel_monkey#/media/File:Saimiri_sciureus-1_Luc_Viatour.jpg)

Marmoset: Carmem A. Busko (https://en.wikipedia.org/wiki/Marmoset#/media/File:Marmoset_copy.jpg)

Chicken: Andrei Niemimäki (https://en.wikipedia.org/wiki/Chicken#/media/File:Male_and_female_chicken_sitting_together.jpg)

Silkie: Benjamint444 (https://en.wikipedia.org/wiki/Silkie#/media/File:Silky_bantam.jpg) Crested partridge: Brian Gratwicke

Common Quail: KaouroV (https://en.wikipedia.org/wiki/Common_quail#/media/File:A_common_quail_in_Lebanon.jpg)

Vulturine guineafowl: Sumeet Moghe (https://en.wikipedia.org/wiki/Vulturine_guineafowl#/media/File:Vulturine_Guineafowl_at_Samburu.jpg)

Black-necked Swan: Sanjay Acharya (https://en.wikipedia.org/wiki/Black-necked_swan#/media/File:Black-necked_Swan.jpg)

Garganey: Dick Daniels (http://theworldbirds.org/)

Mallard: Richard Bartz (https://en.wikipedia.org/wiki/Mallard#/media/File:Anas_platyrhynchos_male_female_quadrat.jpg)

Orangutan: Eleifert (https://en.wikipedia.org/wiki/Orangutan#/media/File:Orang_Utan,_Semenggok_Forest_Reserve,_Sarawak,_Borneo,_Malaysia.JPG)

Green monkey: tjabeljan (https://en.wikipedia.org/wiki/Green_monkey#/media/File:Gambia06Bijilo0015_(5421078756).jpg)

Snub-nosed monkey: Giovanni Mari (https://en.wikipedia.org/wiki/Golden_snub-nosed_monkey#/media/File:Golden_Snub-nosed_Monkeys,_Qinling_Mountains_-_China.jpg)

Baboon: Muhammad Mahdi Karim (https://en.wikipedia.org/wiki/Baboon#/media/File:Olive_baboon_Ngorongoro.jpg)

Macaque: Charles J. Sharp (https://en.wikipedia.org/wiki/Rhesus_macaque#/media/File:Rhesus_macaque_(Macaca_mulatta_mulatta),_male,_Gokarna.jpg)

Ural owl: Jyrki Salmi(https://en.wikipedia.org/wiki/Ural_owl#/media/File:Strix_uralensis,_Kotka,_Finland_1.jpg)

Tawny owl: Martin Mecnarowski (https://en.wikipedia.org/wiki/Tawny_owl#/media/File:Strix_aluco_3_(Martin_Mecnarowski).jpg)

Eurasian eagle owl: Martin Mecnarowski (https://en.wikipedia.org/wiki/Eurasian_eagle-owl#/media/File:Bubo_bubo_3_(Martin_Mecnarowski).jpg)

Long-eared owl: Sascha Rösner (https://en.wikipedia.org/wiki/Long-eared_owl#/media/File:Waldohreule_in_freier_Wildbahn.jpg)

Little owl: Arturo Nikolai (https://en.wikipedia.org/wiki/Little_owl#/media/File:Mochuelo_Com%C3%BAn_(_Athene_noctua_)(1).jpg)

Supplementary Figure 4:

Penaeus monodon: Author unknown (https://commons.wikimedia.org/wiki/File:Penaeus_monodon.jpg)

Common Octopus: Author unknown (https://commons.wikimedia.org/wiki/File:Octopus2.jpg)

Lobster: U.S. National Oceanic and Atmospheric Administration (https://www.flickr.com/photos/noaaphotolib/5114738480/)

Lancelet: Hans Hillewaert (https://upload.wikimedia.org/wikipedia/commons/4/47/Branchiostoma_lanceolatum.jpg)

Acorn worm: NOAA Photo Library (https://en.wikipedia.org/wiki/Acorn_worm#/media/File:Expn7526_(38827990315).jpg)

Lamprey: Bulletin of the United States Fish Commission (https://en.wikipedia.org/wiki/Arctic_lamprey#/media/File:Lampetra_camtschatica.jpg)

Alitta succinea: Hans Hillewaert (https://commons.wikimedia.org/wiki/File:Alitta_succinea_(epitoke).jpg).

## Author contributions

J. K. and C.B. designed the study. J.K. established and annotated the sample collection. A.P., C.Se., M.K., J.S.L.R., L.K., A.E., S.K., A.B., M.L., D.T., P.B., M.H., M.D., D.L.D., and A.K.H. provided samples and sample annotations. J.K., A.N., C.Se., L.C.S., M.C.K., J.S.L.R., L.K., A.E., D.P., S.K., C.Sc., M.F., K.S, and M.D. prepared samples and isolated DNA. A.N. performed DNA methylation profiling with support from P.D. J.K., D.R., and B.E. processed the data. J.K. and D.R. analyzed the data with support from N.F. J.K., D.R., and C.B. interpreted the data. J.K., D.R., and C.B. wrote the manuscript with contributions from all authors.

## Data and materials availability

All data and materials are available via the Supplementary Website (http://cross-species-methylation.computational-epigenetics.org). In addition, the raw and processed data (converted and unconverted RRBS libraries) are available from the NCBI Gene Expression Omnibus (GEO) repository (accession number: <u>GSE195869</u>). The raw BAM files comprise two terabytes and the processed data comprise 550 gigabytes, corresponding to 580 individual species. The source code (R and Python) underlying the current version of the manuscript is available as a ZIP archive on the Supplementary Website (http://cross-species-methylation.computational-epigenetics.org). The source code underlying the final version of the manuscript will be provided as a GitHub repository and as a permanent code archive on Zenodo.

## Materials and Methods

### Sample collection

Samples were selected to represent all vertebrate classes and proximal marine invertebrates. To obtain this breadth of coverage, tissue samples were obtained from several sources (**Supplementary Table 1-4**):

1. *Research Institute of Wildlife Ecology of the University of Veterinary Medicine Vienna* (1619 samples): Tissue samples were collected and frozen during routine pathological examination of deceased wild, pet, and zoo animals. Samples were stored at -80 °C. Pathological conditions and sample preservation (1: well preserved, 2: intermediate, 3: rotten) were recorded (**Supplementary Table 3**). Well-preserved samples were preferentially selected. Species names were obtained from the notes of the pathological examination. Peripheral blood of Bactrian camel (*Camelus bactrianus*) and llama (*Lama glama*) was collected as part of routine veterinary examinations. Blood cell types were isolated using forward/side scatter FACS^44^.
2. *Ocean Genome Legacy Center (OGL) at the Northeastern University Marine Science Center* (602 samples): Specimens were collected and deposited to the OGL collections by numerous collaborating researchers and were stored at -80°C prior to dissection. DNA was isolated using the Qiagen DNeasy Blood & Tissue kits according to the manufacturer’s protocol and stored at -80°C prior to shipment on dry ice.
3. *Commercial fish farm (Biofisch Wien)* (73 samples): Innards of fish killed for food were immediately dissected, transported on dry ice, and stored at -80 °C until DNA extraction using the Qiagen kit.
4. *Commercial fish retailer (Naschmarkt Wien)* (67 samples): Whole specimens of sea food were purchased, transported on ice, dissected, and stored at -80 °C until DNA extraction using the Qiagen kit.
5. *Department of Medical Biochemistry of the Medical University of Vienna* (21 samples): Tissue samples of chickens (*Gallus gallus*) were collected and stored at -80 °C until DNA extraction using the Qiagen kit.
6. *Max Planck Institute for Evolutionary Biology* (16 samples): Tissue samples of Eurasian blackcaps (*Sylvia atricapilla*). The birds were caught at the Pape Ornithological Station, Latvia (56°9′48′′N, 21°1′35′′E) between the end of August and the beginning of September 2011, then transported to the University of Ferrara, Ferrara, Italy, where they were held in aviaries till sample collection. Experimental procedures unrelated to this work were carried out during the autumn migratory season 2011, and birds were stored at - 80 °C until organs were dissected in 2016 at the MPI for Evolutionary Biology. DNA was isolated using a standard phenol-chloroform extraction protocol and stored in ddH2O at -80°C.
7. *Department of Biology of the University of Kentucky* (16 samples): Tissue samples of Mexican axolotls (*Ambystoma mexicanum*) were collected and stored at -80 °C until DNA using a standard phenol-chloroform extraction protocol.
8. *CeMM Research Center for Molecular Medicine of the Austrian Academy of Sciences* (12 tissue samples): Tissue samples of Tasmanian devils (*Sarcophilus harrisii*) (healthy controls) were collected and processed as part of a previously published study that investigated Tasmanian devil transmissible tumors^102^.
9. *St. Anna Children’s Cancer Research Institute* (12 samples): Leukocytes and erythrocytes of zebrafish (*Danio rerio*) were collected from kidneys and blood of adult animals. Cells were dispersed in PBS supplemented with 3% FCS and 2 mM EDTA and sorted by FACS following an established protocol for blood cell populations in zebrafish^103^. Sorted cells were lysed and DNA was isolated using the Qiagen kit.
10. *Department for Pathobiology of the University of Veterinary Medicine Vienna* (7 samples): Tissue samples of flying snakes (*Chrysopelea*) were collected during routine pathological examination of deceased animals and stored at -20 °C until DNA extraction using the Qiagen kit.

### Taxonomic annotation

All samples were annotated with a scientific (Latin) name and a common (English) name based on the information provided by the sample source. Occasionally, the available information did not support the assignment of the exact species; these samples were assigned genus names rather than individual species names. Moreover, the sequencing data for each species were compared with public reference databases (as described in more detail below), and potential errors or ambiguities were flagged or corrected based on manual inspection. Detailed taxonomic annotations for all included species were obtained from the NCBI database using the function *classification* in the R package *taxize* and manually reviewed for accuracy. In all analyses, marsupials were placed in their own group (although they are mammals), given their unique evolutionary history. We used the NCBI Taxonomy Browser (https://www.ncbi.nlm.nih.gov/Taxonomy/CommonTree/wwwcmt.cgi) to create a taxonomic tree for all species, and we visualized the resulting phylogenetic relationships and associated information using the iTOL software^104^. The resulting annotated species tree is provided for interactive viewing and browsing under the following URL: https://itol.embl.de/tree/841339292169571630660457.

### DNA extraction from tissue samples

DNA from tissue samples was extracted using the Qiagen DNeasy Blood & Tissue kits according to the manufacturer’s protocol. Briefly, small pieces of tissue (∼2 mm^3^) were placed in collection tubes, covered with proteinase-K containing digestion buffer, and shaken overnight at 56 °C. When the tissue samples were completely dissolved, the DNA was bound to spin columns and washed, followed by elution in 50-200µl nuclease free water, depending on the expected amount of DNA. The DNA concentration was then quantified using a Qubit fluorometer. DNA from isolated blood cells was isolated as previously described^44^, using the Allprep DNA/RNA Mini kit (Qiagen). Between 50,000 and two million cells were lysed in 600 μl Buffer RLT Plus supplemented with 1% β-Mercaptoethanol and vortexed thoroughly for at least 5 min. The procedure of isolating DNA and RNA was performed according to protocol. DNA was stored at -20 °C.

### DNA methylation profiling by RRBS

Reduced representation bisulfite sequencing (RRBS) was performed as described previously^44, 69^, using 100 ng of genomic DNA for most samples, while occasionally going down to 1 ng for samples with low DNA amounts (Supplementary Table 1). To assess the bisulfite conversion efficiency independent of CpG context, methylated and unmethylated spike-in controls were added at a concentration of 0.1%. For most samples, DNA was digested using the restriction enzymes MspI and TaqI in combination (as opposed to only MspI in the original protocol) in order to increase genome-wide coverage. For certain older samples, only MspI was used (**Supplementary Table 1**). Restriction enzyme digestion was followed by fragment end repair, A-tailing, and adapter ligation. Finally, the libraries were size selected by performing a 0.75× cleanup with AMPure XP beads (Beckman Coulter, A63881) retaining fragments of about 100 bp to 1000 bp length. The amount of effective library was determined by qPCR, and samples were multiplexed in pools of 10 with similar qPCR *Ct* values. The pools were then subjected to bisulfite conversion, followed by library enrichment with PCR. Enrichment cycles were determined using qPCR and ranged from 6 to 18 (median: 11). After confirming adequate fragment size distributions on Bioanalyzer High Sensitivity DNA chips (Agilent), libraries were sequenced on Illumina HiSeq 3000/4000 machines using the 50 or 60 bp single-read setup.

### Sequencing of unconverted RRBS libraries

To distinguish with confidence between genomic thymines und constitutively unmethylated cytosines (which are read as thymines using bisulfite sequencing), we sequenced one RRBS library for each species omitting the bisulfite conversion. Libraries were multiplexed in pools of up to 20 samples, and the pools were subjected to size selection with a 0.6x reverse bead clean up and eluted in 20 µl EB. The amount of effective library in the size-selected pools was determined by qPCR using 1 µl size selected library as input. Based on qPCR *Ct* values for each pool, the number of PCR enrichment cycles was determined as the *Ct* minus two, which ranged from 5 to 11 cycles. PCR and qPCR cycler programs were the same as in the RRBS protocol. The enriched libraries were subjected to a 1.0x bead clean up. Library size distributions were assessed on Bioanalyzer High Sensitivity DNA chips (Agilent) and ranged from 260 to 300 bp (mostly 280 bp). Libraries were sequenced on Illumina HiSeq 3000/4000 machines using the 50 or 60 bp single-read setup.

### RRBS data processing

The RRBS data were processed using an updated version of the RefFreeDMA software^44^, which is available on Github (https://github.com/jklughammer/RefFreeDMA). For each species, we used RefFreeDMA to cluster the sequencing data into read stacks corresponding to specific positions in the genome, to infer the genomic DNA sequence as a weighted consensus for each read stack (including both converted and unconverted RRBS libraries), and to perform DNA methylation calling for each sample against these consensus reference fragments. Two improvements were introduced in the process of generating the consensus references: (i) Detection and removal of contaminating microbial sequences by mapping all reads to a “decoy” genome consisting of the NCBI BLAST dataset of representative bacterial/archeal genomes and keeping only unmapped reads; (ii) incorporation of unconverted RRBS libraries to enhance detection of consistently unmethylated genomic cytosines. For analyses that focused on the genomic sequence (e.g., k-mer frequencies, sequence-based prediction of DNA methylation), only those consensus reference fragments that were covered by the corresponding uncovered RRBS library were considered, in order to minimize bias. Finally, summary statistics and quality metrics (including mapping rate, number of covered CpGs, conversion efficiency, DNA methylation level, contamination level, pre-fragmentation) were calculated for each sample (**Supplementary Table 1**).

### RRBS coverage simulation

To assess the genomic coverage of RRBS across a wide range of species, we simulated the restriction digest and size selection in RRBS for all annotated vertebrate genomes that were available from the UCSC Genome Browser, and we determined the expected RRBS coverage for relevant genomic elements (CpG islands, transcripts, promoters, and repeats) in each species. To create in silico RRBS libraries, we first mapped all MspI and TaqI restriction sites in the corresponding genomes using the *matchPattern* function from the R package *Biostrings*^105^. The resulting restriction fragments were then filtered to mirror the RRBS size selection step, retaining fragments with a length between 50 bp and 1000 bp. Of these fragments, the first and last 50 bp were registered as simulated RRBS reads. We next identified all CpGs within the genomes using the *matchPattern* function and intersected these coordinates with the regions covered by the in silico RRBS libraries, and with the relevant genomic elements (CpG islands, transcripts, promoters, and repeats) using the *findOverlaps* function of the R package *GenomicRanges*^106^. Finally, we calculated the fraction of CpGs within each of the assessed genomic elements that are covered by the in silico RRBS libraries. For each genome, the coordinates of the genomic elements were downloaded from the UCSC Genome Browser website (goldenpath/<GE-nome>/biZips) using the *rtracklayer*^107^ package. Promoters were defined as the regions 1000 bp upstream and 500 bp downstream of the transcription start sites. The genome sequences were obtained from the corresponding genome assemblies provided by the UCSC Genome.

The following species and genome assemblies were included in the analysis: C. intestinalis (ci3), African clawed frog (xenLae2), Armadillo (dasNov3), Elephant shark (calMil1), Tibetan frog (nanPar1), Lizard (anoCar2), Medaka (oryLat2), Fugu (fr3), Tetraodon (tetNig2), Nile tilapia (oreNil2), Kangaroo rat (dipOrd1), Stickleback (gasAcu1), Atlantic cod (gadMor1), Sloth (choHof1), Zebrafish (danRer11), Manatee (triMan1), Microbat (myoLuc2), Mouse (mm39), Garter snake (thaSir1), Naked molerat (hetGla2), Squirrel (speTri2), Zebra finch (taeGut2), Golden eagle (aquChr2), Chinese hamster (criGri1), Guinea pig (cavPor3), S. purpuratus (strPur2), Brown kiwi (aptMan1), Mouse lemur (micMur2), Hawaiian monk seal (neoSch1), Chicken (galGal6), Budgerigar (melUnd1), American alligator (allMis1), Elephant (loxAfr3), Lamprey (petMar3), Turkey (melGal5), Painted turtle (chrPic1), Cow (bosTau9), Ferret (musFur1), Rabbit (oryCun2), Tree shrew (tupBel1), Hedgehog (eriEur2), White rhinoceros (cerSim1), Wallaby (macEug2), Marmoset (calJac4), Sheep (oviAri4), Megabat (pteVam1), Squirrel monkey (saiBol1), Cat (felCat9), Tasmanian devil (sarHar1), Golden snub−nosed monkey (rhiRox1), Pig (susScr11), Rhesus (rheMac10), Baboon (papAnu4), Orangutan (ponAbe3), Alpaca (vicPac2), Horse (equCab3), Green Monkey (chlSab2), Dog (canFam5), Rat (rn7).

Because many reference genomes had an incomplete assembly status and comprised many individual scaffold sequences (often 10,000s or 100,000s thousands instead of a few dozen chromosomes), we concatenated individual sequences into 20 artificial chromosomes, separating the sequences by stretches of 100 Ns. This improved runtimes and avoided out-of-memory issues. After processing, genomic coordinates based on the artificial chromosomes were ported back to the original coordinate space to match the genome annotations.

### Read coverage analysis

To assess biological and technical effects on our RRBS libraries and on the derived consensus references, each consensus reference fragment was evaluated based on its read coverage across all samples for a given species (**Supplementary Figure 2a**). The following classification was applied for each sample: If a fragment had a read coverage of more than half the average coverage in that sample it was considered reliably covered. If a fragment had a coverage of more than four times the average coverage across that sample it was considered highly covered. Next, fragments that were highly covered in more than 80% of the samples were labeled as “Repeat” to indicate that they were likely derived from repetitive genomic regions; fragments that were highly covered in less than 20% of the samples were labeled as “Amplified”, given that this pattern is characteristic of PCR amplification artefacts; and fragments that were reliably covered in more than 80% of the samples of one individual but in less than 20% of the samples of other individuals were labeled “Private”, as such patterns can arise from inter-individual genetic variability. For each sample, the relative proportion of these three categories (“Repeat”, “Amplified”, “Private”) was calculated and averaged across all samples for a given species. For downstream analysis, only species with at least four samples and at least two individuals were considered.

### Cross-mapping analysis

To validate our consensus references and to perform gene-centric analyses, we devised a cross-mapping workflow that connects the RefFreeDMA-derived consensus reference fragments to the most fitting reference genomes. We pursued an empirical “best fit” approach by aligning all consensus reference fragments for a given species to all reference genomes available in the UCSC Genome Browser within the same class (as determined by the *lowest_common* function of the R package *taxize*). For each species, the reference genome with the highest mapping rate was determined and used for further analysis. Mapping was performed using the crossmapping function of RefFreeDMA with an allowed mismatch rate of up to 0.2 (this value was empirically determined). The genomes used for cross-mapping and their preparation are described in the *RRBS coverage simulation* section. DNA methylation profiles across transcripts were created by averaging DNA methylation calls within five kilobases upstream or downstream of the gene body in bins of 100 base pairs, and in bins of 200 base pairs within the gene body itself. For sample-wise analyses the samples were kept separate, whereas all samples of a given species were combined for species-wise analyses.

### Integration of publicly available WGBS data

To validate our RRBS-based, reference-free DNA methylation analysis with publicly available WGBS data, we identified those species in our dataset for which WGBS data were available from GEO, which included: *Phascolarctos cinereus*^57^ (GSE149600), *Bos taurus*^56^ (GSE147087), *Mus musculus*^55^ (GSE42836), *Gallus gallus*^65^ (GSE146620), *Parus major*^64^ (SRR2070790), *Chelydra serpentina*^63^ (data provided by the authors), *Xenopus laevis*^61, 62^ (GSE76247, GSE90898), *Danio rerio*^59, 60^ (GSE149416, GSE134055), *Callorhinchus milii*^25^ (GSE141609), *Branchiostoma lanceolatum*^25, 58^ (GSE102144, GSE141609), *Crassostrea gigas*^54^ (GSE40302), and *Octopus bimaculoides*^25^ (GSE141609). For all species except *Danio rerio* and *Parus major*, supplementary files containing CpG-wise coverage and methylation information were available on GEO and obtained using the GEOquery package^108^. All files were converted into a common format containing CpG-wise read coverage and DNA methylation ratio. Data for *Danio rerio* (GSE149416, GSE134055) and *Parus major* (SRR2070790) were processed starting from the raw sequencing data using the gemBS pipeline^109^ with the danRer11and the Parus_major1.1 assembly, respectively.

### Validation of species annotations

We used the DNA sequencing data to verify species annotations. First, we created a bisulfite converted version of the NCBI BLAST nucleotide database (Nucleotide collection nr/nt), and we mapped 1000 randomly selected reads per sample with the NCBI BLAST command line tool, using the following parameters: -max_target_seqs 100 -num_threads 4 -word_size 15 -evalue 0.00000001 -outfmt “6 qseqid sseqid sscinames scomnames qlen slen sstart send pident length evalue bitscore qseq” (https://github.com/jklughammer/bisulfiteBlast). Where both of the two best matching species differed by more than the level of “class” from the annotated species (this was assessed using the *lowest_common* function of the R package *taxize*), samples were manually inspected and flagged as “unreliable” if the discrepancies could not be explained (e.g., by the absence of related species in the NCBI database). In total, 30 samples were flagged as unreliable (**Supplementary Table 1**).

### Analysis of genome-wide DNA methylation levels

To investigate the association between genome-wide DNA methylation levels and the genomic DNA sequence composition, we calculated the mean DNA methylation levels across all CpGs and samples for each species, and we correlated it with three sets of features derived from the corresponding consensus reference: (i) k-mer frequencies; (ii) CG composition; (iii) CpG island frequencies. K-mer frequencies for k ranging from 1 to 3 were calculated using the MEME suite’s *fasta-get-markov* software tool^110^. CG composition included the frequency of C and G nucleotides, the frequency of CpG dinucleotides, the ratio between observed and expected CpG frequencies (where the expected frequency is defined as the calculated combinatory frequency based on independent C and G frequencies), and the absolute number of covered CpG sites. CpG island frequencies were calculated by determining the percentage of consensus reference fragments that fulfilled the Gardener-Garden and Takai-Jones criteria for CpG islands^111, 112^, requiring a GC content (combined C and G frequencies) of at least 50% (Gardiner-Garden) or 55% (Takai-Jones) and a CpG observed vs. expected ratio of at least 0.6 (Gardiner-Garden) or 0.65 (Takai-Jones), over stretches of 50 bp.

We evaluated the explanatory power of these feature groups for the observed variation in genome-wide DNA methylation levels across species, using a standard linear model as well as linear models that included the phylogenetic group annotation or the taxonomy tree as additional prior features. Linear models were implemented using the R package *phylolm*^113^ to facilitate integration of taxonomy tree structure. The variance explained (R^2^) by each of the models, was calculated using the *R2.pred()* function from the *rr2* package^114^. The R^2^ values were further adjusted using the Wherry Formula-1 formula to account for the number of variables in each of the models^115^. All models were additionally evaluated by the standard Akaike information criterion, using the *AIC* function from the *stats* or *phylolm* package respectively. To evaluate the predictive importance of the different 3-mers, we used stepwise feature selection^116^, iteratively adding and removing the features (individual 3-mers) in the linear model. Models were compared based on the Akaike information criterion (AIC) using the *stepAIC* function from the R package *MASS*. Each 3-mer was assigned a stability score calculated as the percentage of bootstrap experiments in which the feature was selected for the final model.

Finally, we assessed how well 3-mer frequencies recapitulate phylogenetic distance between the analyzed species. To that end, we derived a pairwise distance matrix across species based on the global 3-mer frequencies in each species (calculated across the consensus references), using the *dist* function in R *stats* package^117^. We then performed hierarchical clustering of this distance matrix using the *hclust* function from the same package with default parameters, and we visualized the result as a dendrogram using the dendextend package in R^118^. For the analysis of publicly available WGBS data, genome-wide DNA methylation values were calculated for each sample by averaging across the DNA methylation levels of all CpGs with coverage exceeding five reads.

### Generalized linear models controlling for phylogenetic relationships

To test for association between genome-wide DNA methylation levels and 3-mer frequencies (or other factors such as theoretical cancer risk), we used generalized linear models that explicitly account and control for phylogenetic relationships. Models were built individually for each factor, either with and without taking phylogeny into account, and the corresponding coefficients and associated p-values were used for interpretation. The phylogenetic models were built with the *compare.gee* function from the *ape*^119^ package assuming a gaussian distributions and using the taxonomic tree as depicted in Figure 1c. The standard models (without controlling for phylogeny) were built using the *glm* function from the *stats* package. For the 3-mer analysis, the p-values obtained from both models were adjusted for multiple testing using the Bonferroni method.

### Analysis of DNA methylation erosion

As a measure of DNA methylation erosion, the proportion of discordant reads (PDR) was calculated as described in the original publication^67^. A custom Python script (integrated in RefFreeDMA) was used to determine the number of concordantly or discordantly methylated reads with at least four valid CpG measurements for each CpG within each sample. For each CpG, the PDR was then calculated as the ratio of discordant reads compared to all valid reads covering that CpG. CpGs at the end of a read were disregarded as potentially unreliable. Finally, sample-wise PDR values were calculated by averaging across their CpG-wise values.

### Prediction of locus-specific DNA methylation levels

To investigate the association between locus-specific DNA methylation and the underlying DNA sequence, we trained machine learning classifiers to predict the mean DNA methylation levels of individual genomic regions (averaged across samples and/or tissues in a given species) based on their genomic DNA sequence. Specifically, we trained support vector machines (SVMs) with a spectrum kernel from the R package *kebabs*^120^ to predict the discretized DNA methylation states of consensus reference fragments (low: DNA methylation less than 20% in all samples, high: DNA methylation greater than 80% in all samples, mean coverage greater than 10 reads) based on the DNA sequence composition of the consensus reference fragment. From the set of sequences assigned to high and low DNA methylation state, we randomly selected class-balanced training and test sets comprising 2000 sequences each. In those species where one class contained less than 2000 sequences, the number of sequences for the other class was reduced accordingly, in order to avoid class imbalance.

For each species, training and test set sequences were transformed into feature matrices comprising all k-mer frequencies with a fixed length k. Based on the training set, the optimal regularization value (C) as well as the optimal k-mer length (k) were selected by grid search across C values of 0.01, 0.1, 1, and 10, and across k values from 1 to 10, using 10-fold cross-validation (**Supplementary Figure 6b**). Finally, SVMs were retrained on the complete training set (without cross-validation) using the optimal parameters and evaluated on the test set. Generation of feature matrices, grid search, and model fitting was done using the *kebabs* package in R^120^.

A second set of models was trained and evaluated in each species using only 3-mers. To quantify the predictive power of individual 3-mers in each taxonomic group, we calculated mean feature weights for each 3-mer across all species in that group. These mean feature weights were used to generate sequence logos with the *ggseqlogo* package in R^121^, separately for positive and negative feature weights. The significance of differences in the mean feature weights was assessed using the Wilcoxon rank-sum test (*wilcox.test* function from the *stats* package in R). For each taxonomic group, the top-10 k-mers with the most significant differential feature weights were reported (**Supplementary Figure 6e**). Finally, to test the robustness of these results, all predictions were repeated with less stringent thresholds that include sequences with low DNA methylation levels in any sample as opposed to all samples (low: DNA methylation less than 20% in any sample, high: DNA methylation greater than 80% in all samples), and the ROC-AUC values were compared (**Supplementary Figure 6a**).

For validation based on publicly available WGBS data, we selected those species for which a suitable reference genome and WGBS data with at least two biological replicates were available. The following species and datasets were included: *Bos taurus*^56^ (GSE147087, bosTau9), *Mus musculus*^55^ (GSE42836, mm9), *Gallus gallus*^65^ (GSE146620, galGal5), *Xenopus laevis*^61, 62^ (GSE76247, GSE90898, Xla.v91), *Chelydra serpentina*^63^ (data provided by the authors, ASM1885937v1), *Phascolarctos cinereus*^57^ (GSE149600, phaCin_unsw_v4.1), *Danio Rerio*^59^ (GSE134055, danRer11), and *Branchiostoma lanceolatum*^25, 58^ (GSE102144 and GSE141609, Bl71nemr). All genome assemblies were processed using the Biostrings^105^ package and split in 50 basepair tiles, mimicking the consensus reference fragments. Each tile was annotated with its mean DNA methylation level calculated as the coverage-weighted mean of DNA methylation values for each CpG in the tile. As in the RRBS-based analysis, only sequences with a mean coverage of at least 10 reads across all samples were retained. Sequences with DNA methylation levels greater than 80% in all samples were labeled ‘highly methylated’, whereas those with DNA methylation levels less than 20% in all samples were labeled ‘lowly methylated’. The support vector machine was optimized, trained, and assessed on a balanced subset of 2000 randomly chosen sequences, while ensuring that test and training sequences did not belong to the same chromosome. This procedure was repeated three times in each species, in order to assess the stability of the results.

### Cross-species predictions and inverted species

To assess the generalizability of locus-specific prediction across species, models were trained in one species and tested (without re-training) in a different species. Model performance in each scenario was quantified by receiver operating characteristic area under curve (ROC-AUC) values in unseen test sets of the target species.

These cross-species predictions unexpectedly resulted in a few cases (13 out of 580 species) in which the observed cross-species prediction performance was systematically lower than expected by chance. We referred to those outliers as inverted species, given that the relationship between locus-specific DNA methylation and the underlying DNA sequence appeared to be inverted compared to most other vertebrate and invertebrate species. Specifically, species with average ROC-AUC values below 0.45 compared to all other species were denoted as inverted species, and the taxonomic group with most inverted species (*actinopteri*) was investigated further. To that end, we identified those 3-mers whose feature weights deviated most strongly in the inverted species, as compared to all other species in the same taxonomic group. To test the hypothesis that the observed inversion in the relationship between DNA methylation and DNA sequence was likely caused by recent expansion of heavily methylated repeats in the inverted species, we used the identified 3-mers for repeat identification, calculating the frequencies of 3-mer derived 9-mer repeats (e.g., ACGACGACG) across all consensus reference fragments with high (greater than 80%) and low (less than 20%) average DNA methylation levels.

### Analysis of tissue-specific DNA methylation

To assess the prevalence of tissue-specific versus inter-individual differences in DNA methylation, we focused on species with at least two individuals, two tissues, and one common tissue between individuals, after removing species that had less than 50% average CpG overlap between samples or were flagged in the validation of species annotations. For each of the selected species, we calculated the variance explained by tissue and by individual as the average squared Pearson correlation (R^2^) for the mean DNA methylation levels of the consensus reference fragments across samples. The Pearson correlation was calculated using the *cor* function of the package *stats* in R. The significance of the difference between the variance explained by tissue and by individual between taxonomic groups was calculated using a two-sided paired Wilcoxon test *(wilcox.test* in the package *stats*). Word clouds representing the relative frequency of tissues contributing to the analysis were produced using the function *wordcloud* of the package *wordcloud* in R.

Differentially methylated consensus reference fragments between tissue types (specifically heart and liver) were mapped with RefFreeDMA as described previously^44^. First, differentially methylated CpGs were identified using the R package *limma*^122^, with multiple-testing correction using the Benjamini-Hochberg method. Second, the p-values for individual CpGs within the same consensus reference fragment were combined using an adjusted version of the Fisher’s method^123^. Third, to identify the top-500 most hypermethylated consensus reference fragments in each tissue compared to the other tissue, we used a combined rank approach based on p-value, relative difference, and absolute difference in DNA methylation. Fragments were further required to have a p-value less than 0.05 and an average coverage of at least two reads in both tissues.

### Transcription factor binding site analysis

To identify enriched transcription factor binding motifs among the differentially methylated consensus reference fragments, we tested the binding position-weight matrixes (PWMs) from the 2020 version of the JASPAR database^124^ using the AME tool from the MEME package with default parameters^125^. We scored each motif for enrichment among the top-500 hypermethylated fragments relative to the top-500 hypomethylated fragments, and vice versa. Motifs with multiple-testing adjusted p-values lower that 0.05 were considered significantly enriched. Transcription factors were additionally annotated based on their gene expression levels in human tissues, using the consensus transcript expression levels from the Human Protein Atlas (https://www.proteinatlas.org/about/download). Only transcription factors that have normalized RNA expression values greater than one in heart or liver samples were included in the analysis. Moreover, to explore the tissue specificity of transcription factor binding, we clustered the corresponding transcription factors based on their motif enrichment in heart and liver using hierarchical clustering within the *pheatmap* package^126^. GO term annotation of the selected transcription factors was performed using the GOnet^127^ web-tool (https://tools.dice-database.org/GOnet/) with a custom set of relevant GO terms (search terms “heart”, “liver”, ”hypoxia”, ”detoxification”, ”fluid shear stress”, “glucagon”, “secretion”, “differentiation”, ”regeneration” “cell cycle”, ”glucose homeostasis”, ”thyroid hormone”,” nitrogen compound metabolic process”). The resulting network was downloaded as a json file and visualized using Cytoscape^128^. For better visualization, connections between GO terms were cut and redundant annotations removed. Four transcription factors (ZNF740, KLF9, ZNF263, ZNF384) were not part of any “Biological Processes” GO term and one (KLF16) only of the very broad term “signal transduction”. These factors were excluded from the network.

Transcription factors were annotated by their binding preferences (methylated or non-methylated binding site) based on HT-SELEX experiments for their human homologs^85^. Transcription factors annotated as preferring binding to non-methylated sites whose binding sites were hypomethylated in liver (compared to heart) were classified as “active in liver”. Similarly, transcription factors annotated as preferring binding to methylated sites whose binding sites were hypermethylated in liver (compared to heart) were also classified as “active in liver”. Transcription factors showing the opposite characteristics were labeled “active in heart”. Using a manually curated database of human gene-regulatory networks^86^, we identified the potential targets of these transcription factors and visualized the resulting network using Cytoscape^128^.

### Reference-based analysis

Based on cross-mapping as described above, promoter methylation levels for each annotated and covered gene were calculated as mean DNA methylation levels in the proximal 50 bins of the 5 kilobases upstream regions, corresponding to 2.5 kilobases upstream of the transcription start site. Gene identifiers across all genomes were then annotated by the human homolog; first, the original refSeq IDs were converted to NCBI IDs using the NCBI gene2refseq dictionary; second, NCBI IDs were matched to their human orthologs using the NCBI gene_orthologs dictionary. Both dictionaries were obtained from https://ftp.ncbi.nlm.nih.gov/gene/DATA/.

Dimensionality reduced representations of genes or samples in human homolog space across all samples and species were produced by first filtering the gene/sample matrix, only keeping genes that were covered in greater than 100 samples and samples that covered great than 400 genes (for the sample-wise representation), and genes that were covered in great than 400 samples and samples that covered greater than 50 genes (for the gene-wise representation). This filtering strategy was optimized to produce missing-value-free sample-sample or gene-gene correlation matrices using pairwise complete observations Person’s correlation was calculated using the *cor* function of the R package *stats*. Uniform Manifold Approximation and Projection (UMAP) was then performed on these correlation matrixed using the *umap* function of the R package *uwot* with the following relevant parameters: n_neighbors = 20, min_dist=2, spread=1 (for the sample-wise representation) and n_neighbors = 15, min_dist=0.05, spread=1.5 (for the sample-wise representation). Leiden clustering for the gene-wise UMAP representation was performed using the *cluster_leiden* function from the R package *igraph* with a resolution of 0.06. Gene enrichment analysis on the genes in each Leiden cluster was performed with the *gost* function of the R package *gprofiler2* using GO Biological Processes, GO Molecular Function, and GO Cellular Compartment as databases and an FDR-corrected p-value of 0.05 as significance threshold.

To predict sample properties (e.g., tissue type or evolutionary class) based on promoter methylation levels across species, we exploited the common human ortholog gene-space described above. We used a random forest classifier that is able to handle missing values, as implemented in the *rpart* function of the *rpart* R package. We required each gene to be covered in at least 60% of the samples and each samples to be covered by at least 12% of the genes in the assessed subsets of the data. We focused our analysis on mammals and birds as the most highly represented taxonomic groups, and on liver and heart as the most highly represented tissue types. We performed prediction of tissue type (heart versus liver) and prediction of evolutionary class (mammal vs. bird), separately for mammal and bird species or heart and liver tissues, allowing us to compare predictive performance. 80 (tissue prediction) and 100 (class prediction) species were randomly selected for training the model, and the remaining species were used for testing the model. We split by species (rather than by sample) to make sure that training and test data did not contain samples of the same species, thereby focusing the analyses on cross-species prediction. The procedure was repeated in 100 iterations, where each iteration recorded classification success as well as feature weights. The performance (ROC-AUC values) for each classification task was evaluated, and average feature weights were calculated across all 100 iterations. As a control analysis, the same procedure was applied to scrambled data matrices derived from the actual data matrices by randomly distributing the non-missing values across the non-missing value positions; this analysis maintained the missing-value structure to ensure that this did not contribute to the observed predictions.

### Statistical reporting

Boxplots are specified as follows: center line, median; box limits, upper and lower quartiles; whiskers, 1.5x interquartile range; points, outliers (standard *geom_boxplot* configuration). Error bars represent the standard deviation if not specified differently.

---

**Supplementary Figure 1.**
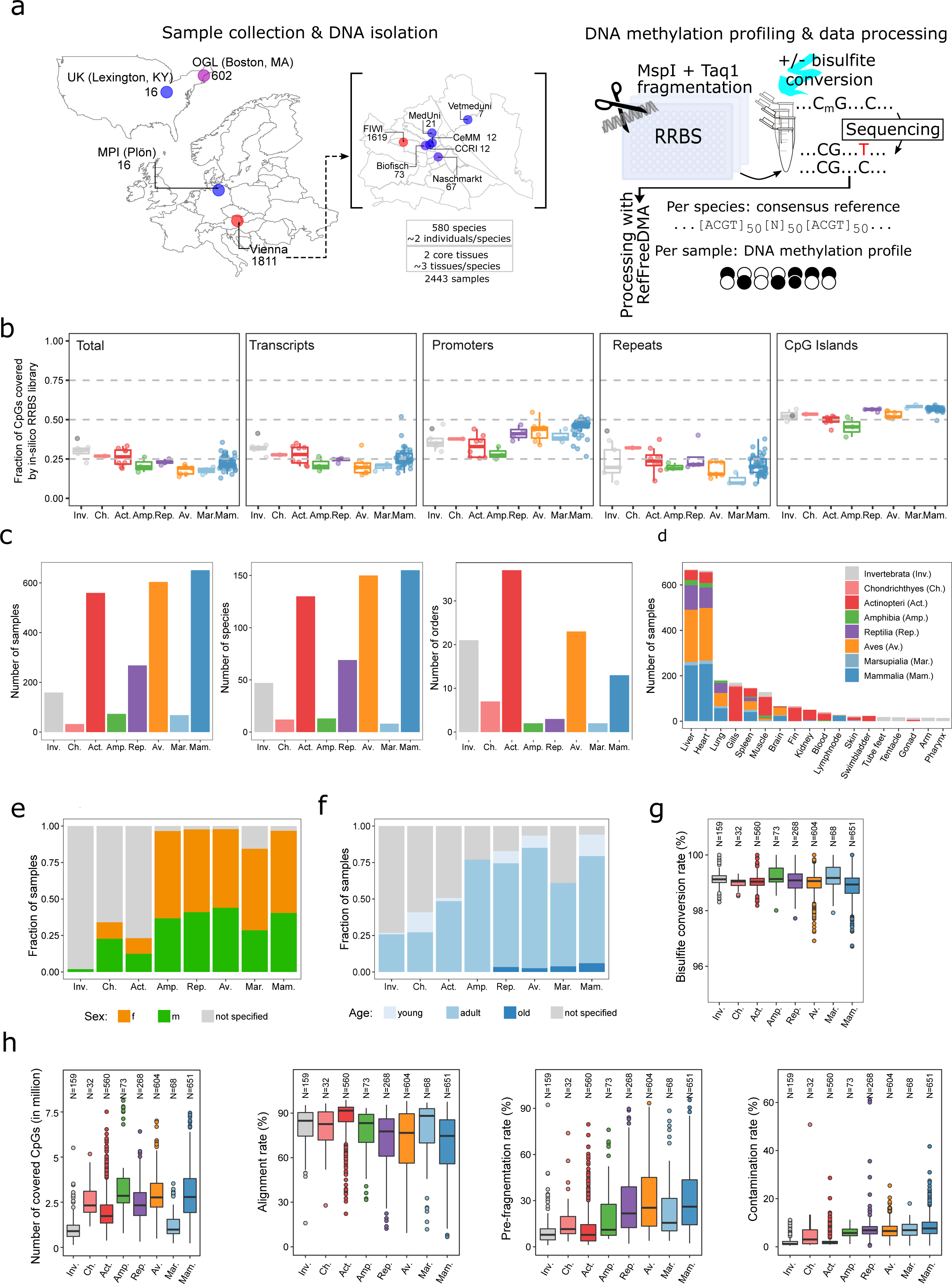
Summary statistics for 2445 DNA methylation profiles from 583 animal species. (a) Schematic overview of the steps taken to assemble the presented resource of vertebrate and invertebrate DNA methylation profiles: Sample collection, DNA isolation, DNA methylation sequencing using the reduced representation bisulfite sequencing (RRBS) assay, and bioinformatic processing using the RefFreeDMA workflow. For each species, an unconverted RRBS library was additionally sequenced to support a more accurate consensus reference reconstruction. Sample sources: FIWI: Research Institute of Wildlife Ecology of the University of Veterinary Medicine Vienna; OGL: Ocean Genome Legacy Center; Biofisch: Commercial fish farm; Naschmarkt: Commercial fish retailer; MedUni: Department of Medical Biochemistry of the Medical University of Vienna; MPI Plön: Max Planck Institute for Evolutionary Biology; UK: Department of Biology of the University of Kentucky; CeMM: CeMM Research Center for Molecular Medicine of the Austrian Academy of Sciences; CCRI: Children’s Cancer Research Institute; Vetmeduni: University of Veterinary Medicine. (b) Boxplots overlayed with individual dots corresponding to individual species, showing the percentage of CpGs in total and within relevant genomic elements (transcripts, promoters, repeats and CpG islands) expected to be covered by RRBS in the corresponding species based on simulations, aggregated by taxonomic groups. (c) Bar plots showing the number of analyzed samples, species, and orders across all taxonomic groups. (d) Bar plot showing the representation of different tissue samples across taxonomic groups. Only tissues with more than ten samples are shown. (e) Stacked bar plot showing the distribution of sex across samples across taxonomic groups. (f) Stacked bar plot showing the distribution of age across samples across taxonomic groups. (g) Boxplots showing the bisulfite conversion efficiency per sample across taxonomic groups. For each sample the higher of two measured values (conversion rate at cytosines outside of a genomic CpG context; conversion rate of unmethylated spike-in controls in the RRBS experiment) is displayed. (h) Boxplots showing RRBS quality control metrics (number of covered CpGs, mapping efficiency, DNA pre-fragmentation, microbial contamination rate) per sample across taxonomic groups.

**Supplementary Figure 2.**
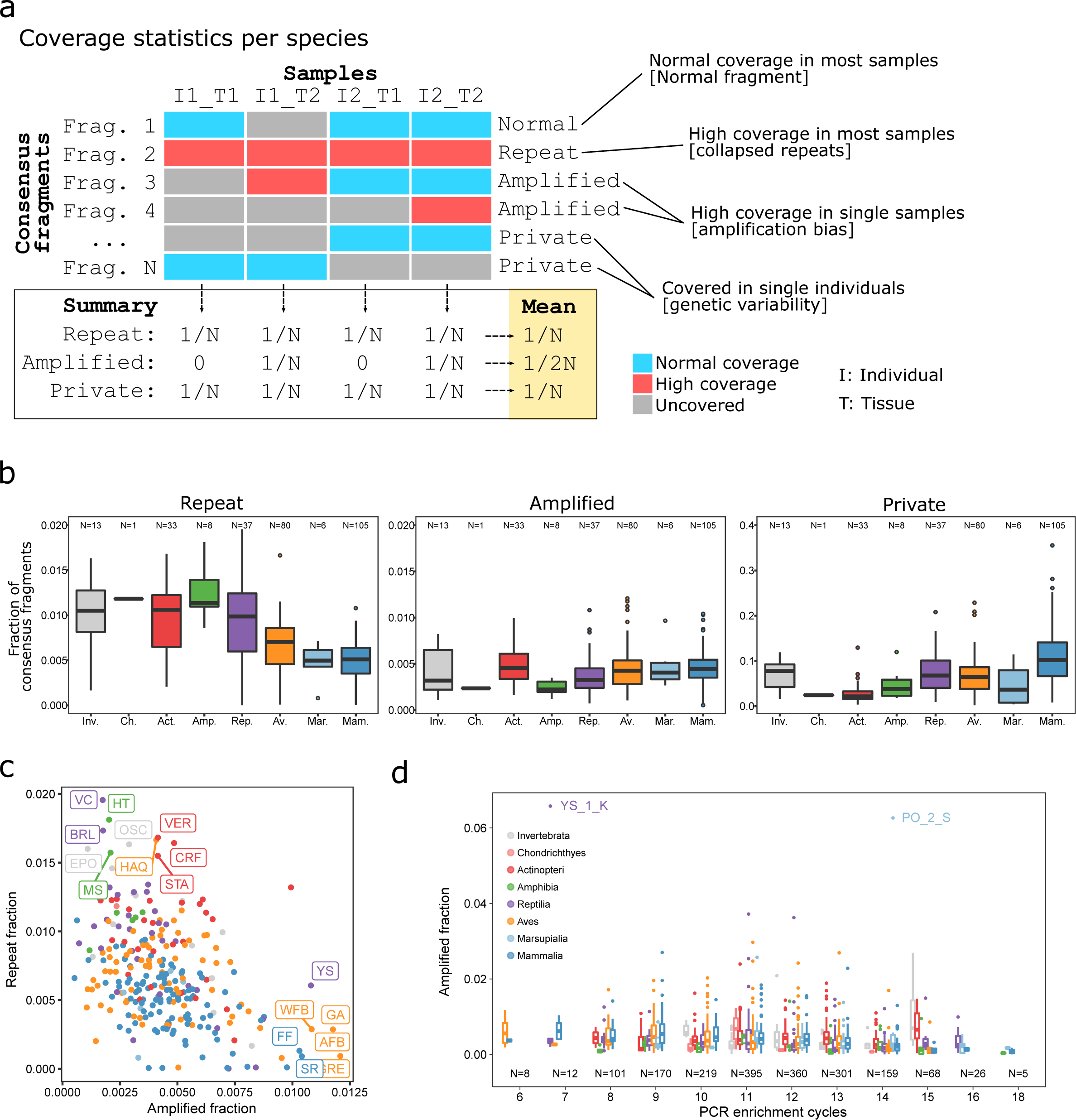
Read coverage analysis for potential technical and biological sources of variability among the consensus references. (a) Schematic overview for the classification of consensus reference fragments as “repeat”, “amplified”, or “private”, and for the calculation of these frequencies within each species. Consensus fragments are classified based on read coverage across all samples in the corresponding species. Sample-wise frequencies of the different classes are calculated, which are then averaged across all samples to generate species-wise measures. (b) Boxplot showing the fraction of consensus fragments for each of the three coverage classes (“repeat”, “amplified”, “private”) in each of the consensus references as defined in (a), aggregated by taxonomic group. (c) Scatterplot showing the relationship between the fraction of consensus fragments classified as “repeat” and those classified as “amplified” for each of the consensus references, colored by taxonomic group. Species at the extremes are annotated with their abbreviations (**Supplementary Table 2**). (d) Boxplot showing the fraction of consensus reference fragments classified as “amplified” within each sample, organized by PCR enrichment cycles and aggregated by taxonomic group. Extreme outliers are annotated with their sample identifiers (**Supplementary Table 1**).

**Supplementary Figure 3.**
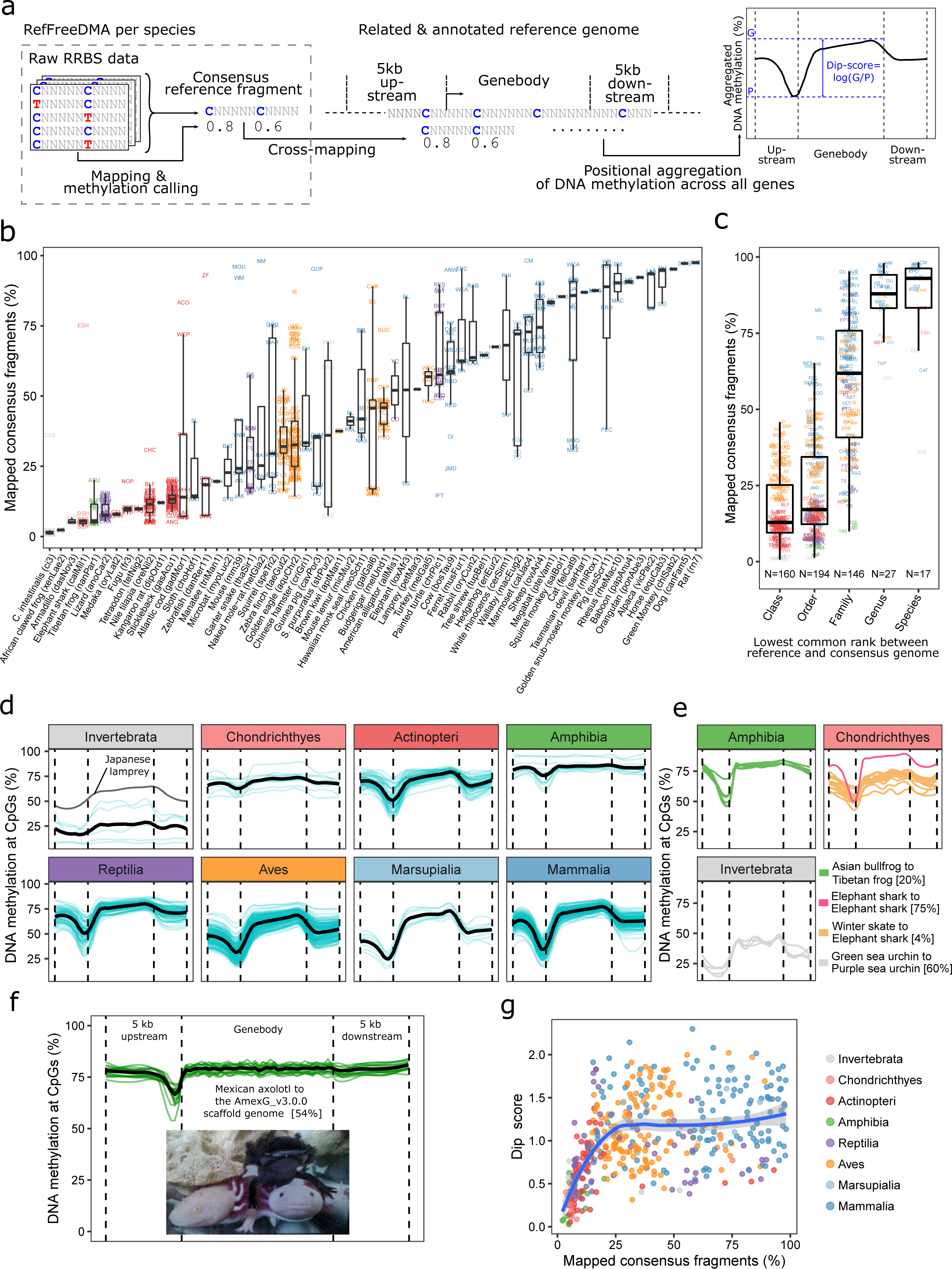
Cross-mapping of consensus reference fragments to gene-annotated reference genomes and analysis of gene-linked DNA methylation patterns. (a) Schematic overview of the cross-mapping approach, which uses gene annotations of related reference genomes to analyze gene-linked DNA methylation patterns including the typical “dip” in promoter regions. Thi is based on RefFreeDMA-derived consensus reference fragments and their DNA methylation levels, which are cross-mapped to gene-annotated reference genomes. The “dip” in DNA methylation at the promoter region is quantified as the log-ratio of DNA methylation levels at gene bodies (G) and at gene promoters (P). (b) Boxplot overlayed by individual datapoints represented as species abbreviations (**Supplementary Table 2**), showing the mapping rates of all consensus references to their best-matching reference genomes (x-axis), colored by taxonomic group. (c) Boxplot overlayed by individual datapoints represented as species abbreviations, showing mapping rates for all consensus references, aggregated by approximated lowest common rank between consensus reference species and reference genome species, colored by taxonomic group. (d) Aggregated and smoothed (using loess with span 0.3) DNA methylation profiles across gene annotations including 5 kilobases upstream and downstream flanking regions. Each thin line represents one species and thick lines represent the average across all species in the respective taxonomic group. (e) Same as (d), displaying individual samples for selected taxonomic groups with less clear profiles. (f) Same as (d) but with less smoothing (using loess with span 0.03), displaying individual samples for Mexican axolotl using an earlier scaffold genome assembly because high-quality gene annotations were not available for the most recent chromosome-scale genome assembly of the axolotl. (g) Scatterplot showing the relationship between consensus fragment mapping rate and dip score for species with at least 4000 gene-associated DNA methylation values, colored by taxonomic group. A loess regression curve fitted to the shown data points is overlaid.

**Supplementary Figure 4.**
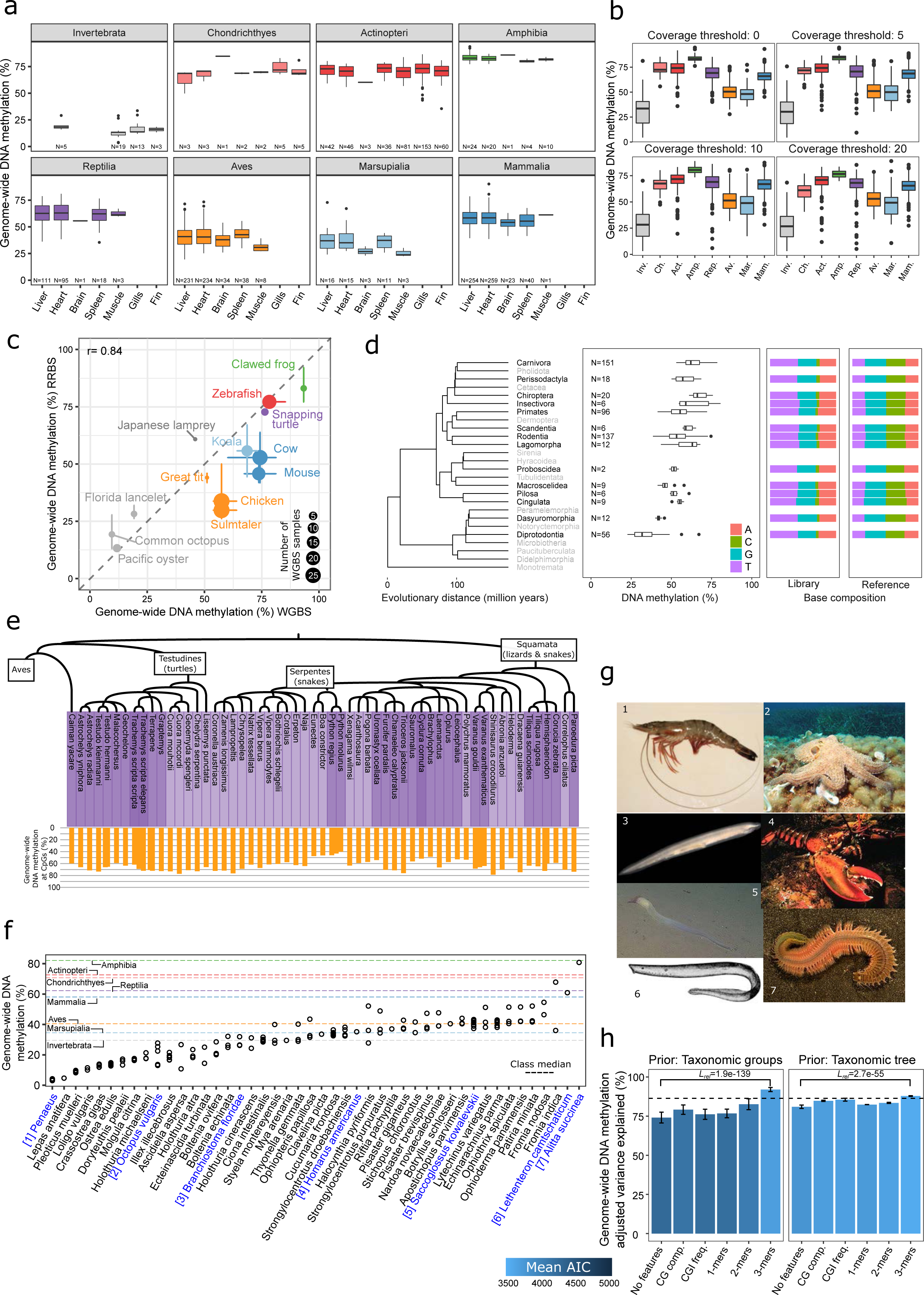
Comparative analysis of genome-wide DNA methylation levels for each species. (a) Boxplots showing mean DNA methylation levels aggregated by taxonomic group and by tissue, including the seven most abundant tissues. (b) Boxplots showing mean DNA methylation levels per sample for different read coverage thresholds, aggregated by taxonomic group. Only CpGs covered by more than the indicated number of reads were included. (c) Scatterplot comparing mean genome-wide DNA methylation levels estimated based on RRBS and WGBS data for twelve species. The dot size indicates the number of samples available for WGBS (range: 1 to 42). Error bars depict minimum and maximum sample-wise values in the respective species and assay. The Pearson correlation (r) between the mean genome-wide DNA methylation levels for RRBS and WGBS is indicated. (d) Boxplot showing mean DNA methylation levels across all mammalian orders, including the marsupial orders diprotodontia (Australian marsupials, mostly herbivores) and dasyuromorphia (Australian carnivorous marsupials). Overlaid are stacked bar plots of base frequencies for the RRBS libraries as well as the consensus reference fragments, indicating broadly similar base frequencies across all mammalian orders. (e) Barplot showing genome-wide DNA methylation levels across all reptilian species ordered by phylogenetic relationships. (f) Scatterplot showing genome-wide DNA methylation levels for individual samples across all invertebrate species as well as Japanese lamprey (*Lethenteron camtschaticum*). The median of genome-wide DNA methylation levels for all taxonomic groups are indicated as dashed lines for reference. (g) Images depicting selected species from panel f. (h) Bar plots showing the percentage of variance explained by features sets reflecting genomic sequence composition (as in panel e), based on linear models that incorporated the taxonomic tree (left) or the taxonomic groups (right) as additional information / priors. All values were adjusted for model complexity (i.e., number of variables) and the colors indicate the mean Akaike information criterion (AIC).

**Supplementary Figure 5.**
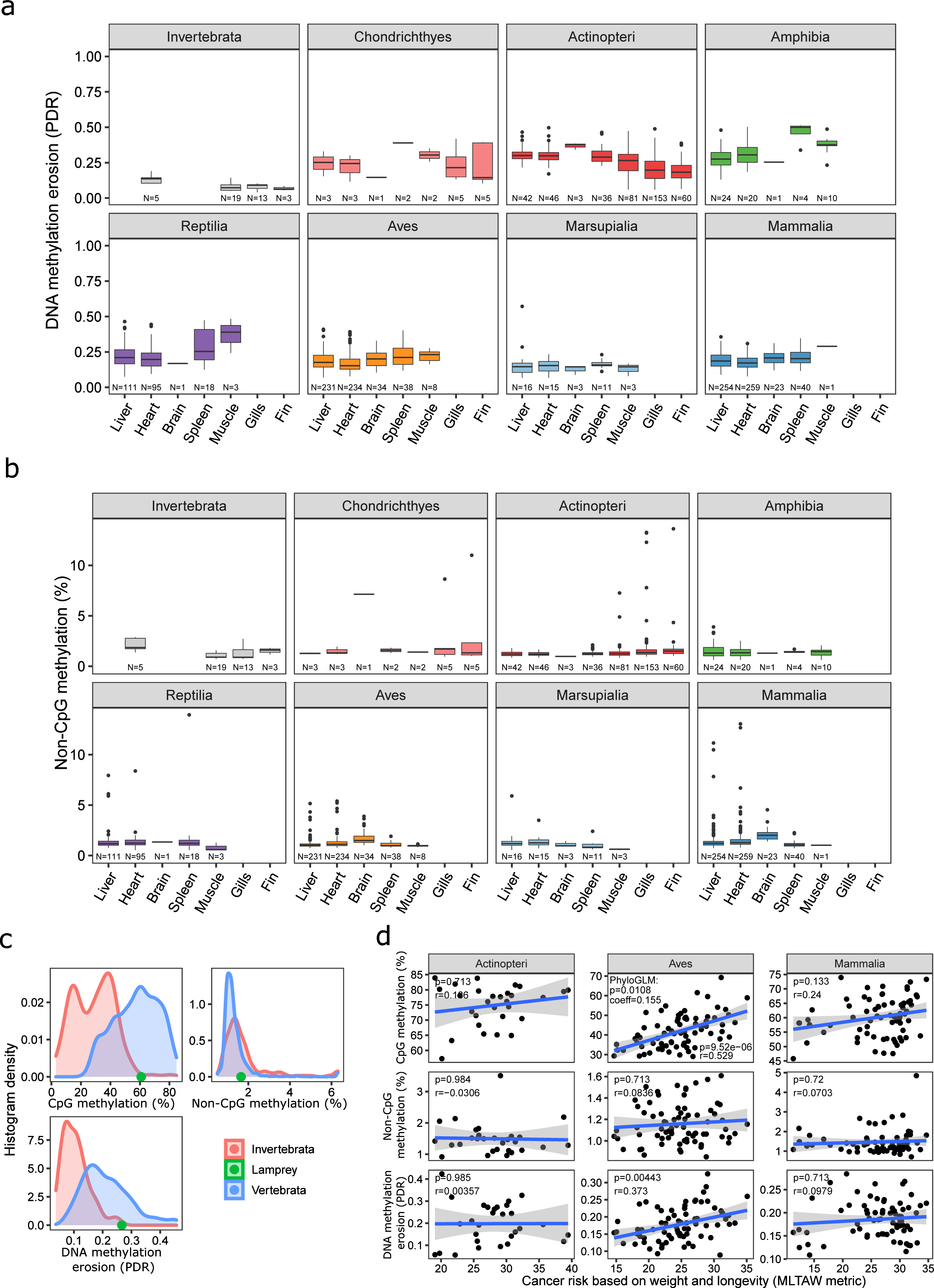
Assessment of non-CpG methylation and DNA methylation erosion as measured by the proportion of discordant reads (PDR) (a) Boxplot showing mean proportions of discordant reads (PDR) as a measure of DNA methylation erosion, aggregated by taxonomic groups and tissues. The seven most frequent tissues are shown. (b) Boxplot showing mean non-CpG methylation levels, aggregated by taxonomic groups and tissues. The seven most frequent tissues are shown. (c) Histograms of genome-wide DNA methylation, non-CpG methylation, and DNA methylation erosion (measured by PDR) for vertebrates and invertebrates to compare to lamprey as a jawless vertebrate (represented as dot). (d) Scatterplot relating genome-wide DNA methylation levels, non-CpG methylation levels, and DNA methylation erosion (measured by PDR) with theoretical cancer risk estimated by the MLTAW metric^77^, which is calculated as log(*Maximum longevity (years)*^6 * *Adult weight (g)*. Pearson’s correlation coefficient and the corresponding significance are indicated. A linear regression curve with a 0.95 confidence interval is overlaid.

**Supplementary Figure 6.**
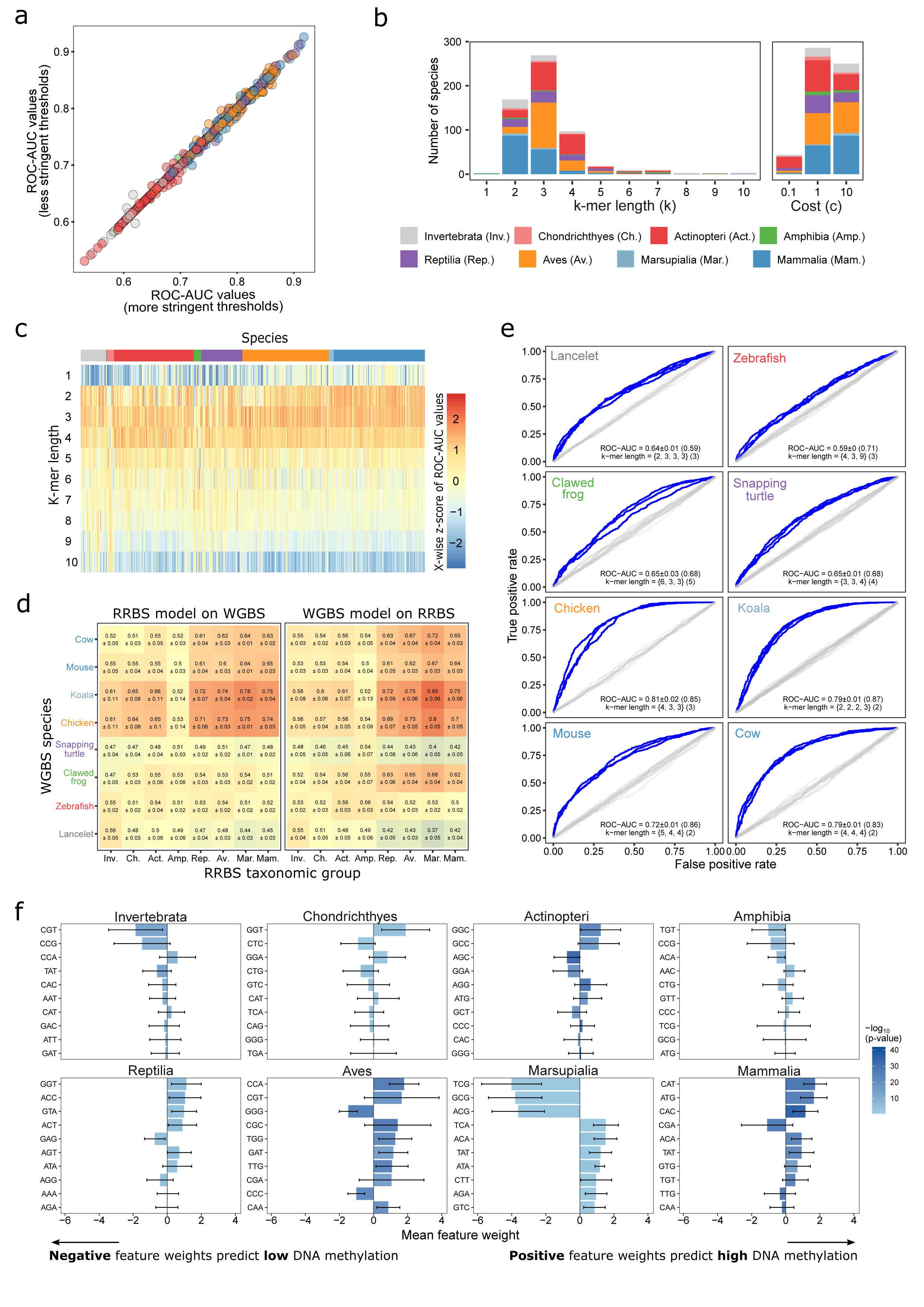
Prediction of locus-specific DNA methylation based on the underlying genomic DNA sequence. (a) Scatterplot comparing the effect of two alternative definitions of highly methylated fragments (DNA methylation levels above 80% in all samples) and lowly methylated fragments (DNA methylation levels below 20% in any sample, x-axis, or in all samples, y-axis) on prediction accuracy measured by ROC-AUC values. Each dot corresponds to one species, colored by taxonomic group. (b) Stacked bar plots displaying the number of species (colored by taxonomic group) for which each k-mer length or learning cost parameter was identified as the optimal one through a grid search on training data. (c) Heatmap showing scaled ROC-AUC values for a range of k-mer lengths (1-10) for prediction of locus-specific DNA methylation across all species, colored by taxonomic group. (d) Heatmap showing the average and standard deviation for ROC-AUC values for locus-specific DNA methylation prediction when training models based on RRBS data and testing with WGBS data, and vice versa. For each taxonomic group, one species with publicly available WGBS was identified and used in this analysis. For each species, the RRBS data of all available samples were combined. (e) ROC curves for prediction of DNA methylation based on the underlying genomic DNA sequence using WGBS data for each of the eight indicated species. Three separate (replicate) ROC curves are shown based on three non-overlapping sets of sequences (blue). ROC-AUC values (mean ± standard deviation) as well as favored k-mer lengths are indicated, and the corresponding values for RRBS-based DNA methylation data are shown in brackets. As negative controls, ROC curves trained and evaluated on data with randomly shuffled labels fall close to the diagonal (in grey). (f) Bar plots displaying the average model feature weight for the 10 most differentially weighted 3-mers across taxonomic groups. Error bars denote standard deviations across all species in the respective taxonomic groups.

**Supplementary Figure 7.**
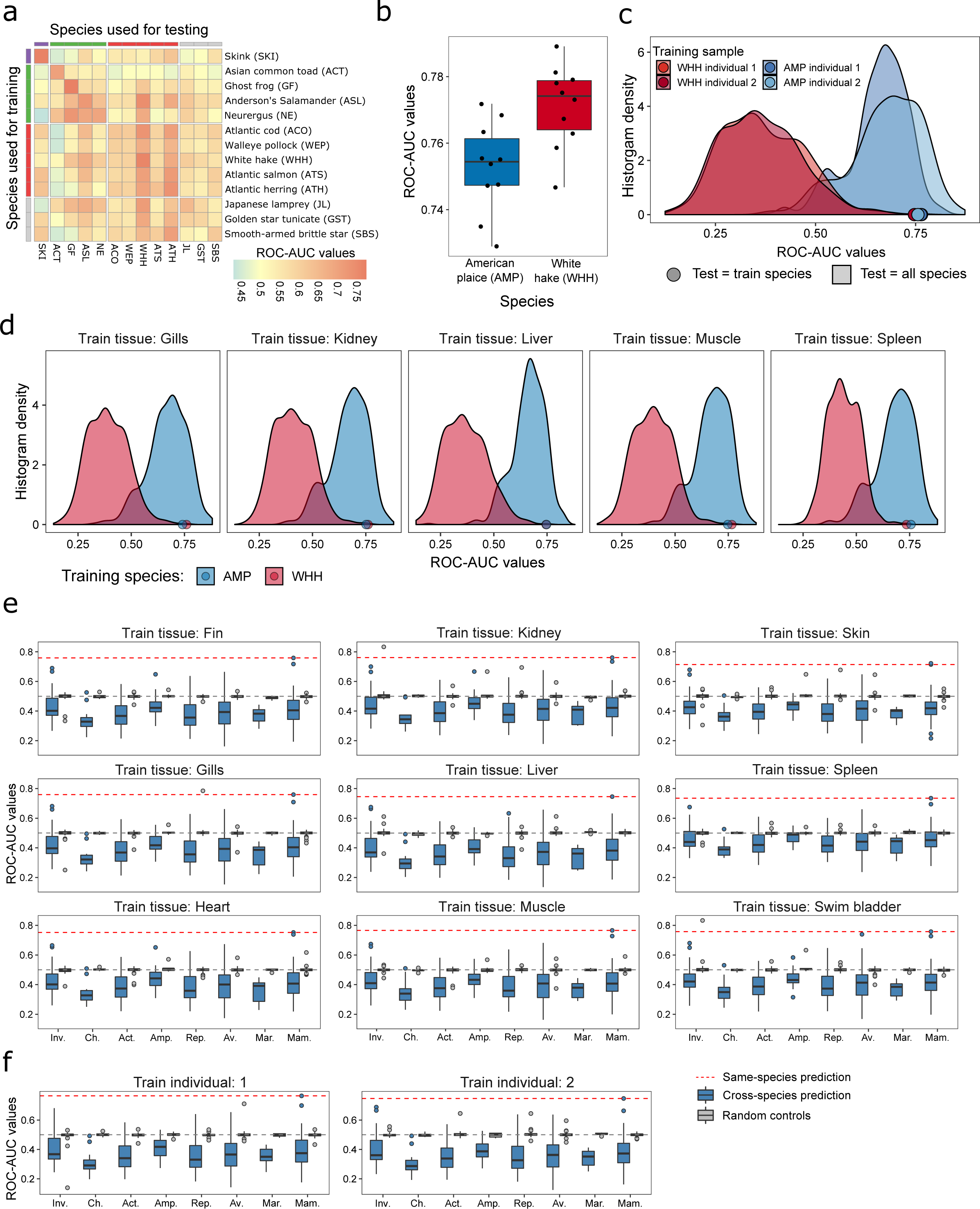
Analysis of inverted species for prediction of locus-specific DNA methylation. (a) Heatmap of cross-species prediction performance (ROC-AUC values), displaying only the inverted species. (b) Bootstrapping stability (ten different selections of training and test data) for the prediction of locus-specific DNA methylation. The training and testing was performed in white hake (WHH, left), a fish species with the inverted pattern, as well as in to American plaice (AMP), a fish species with the non-inverted pattern. (c, d) Histograms of cross-species prediction performance (ROC-AUC values) for an inverted species (white hake, WHH, red) in comparison to one representative non-inverted species (American plaice, AMP, blue). Models were trained and tested separately for each individual (c) and for each tissue (d). Models trained in an inverted species obtained ROC-AUC values below 0.5 in most other species, while models trained in a non-inverted species obtained ROC-AUC values above 0.5 in most other species. (e) Boxplots showing cross-species prediction performance (ROC-AUC values) across all species, using models that were trained on one specific tissue of the inverted species (white hake, WHH). Dashed red lines indicate the ROC-AUC value using the test set from the same species and tissue. (f) Boxplots showing cross-species prediction performance (ROC-AUC values) across all species, using models that were trained on one specific individual of the inverted species (white hake, WHH). Dashed red lines indicate the ROC-AUC value using the test set from the same species and individual.

**Supplementary Figure 8.**
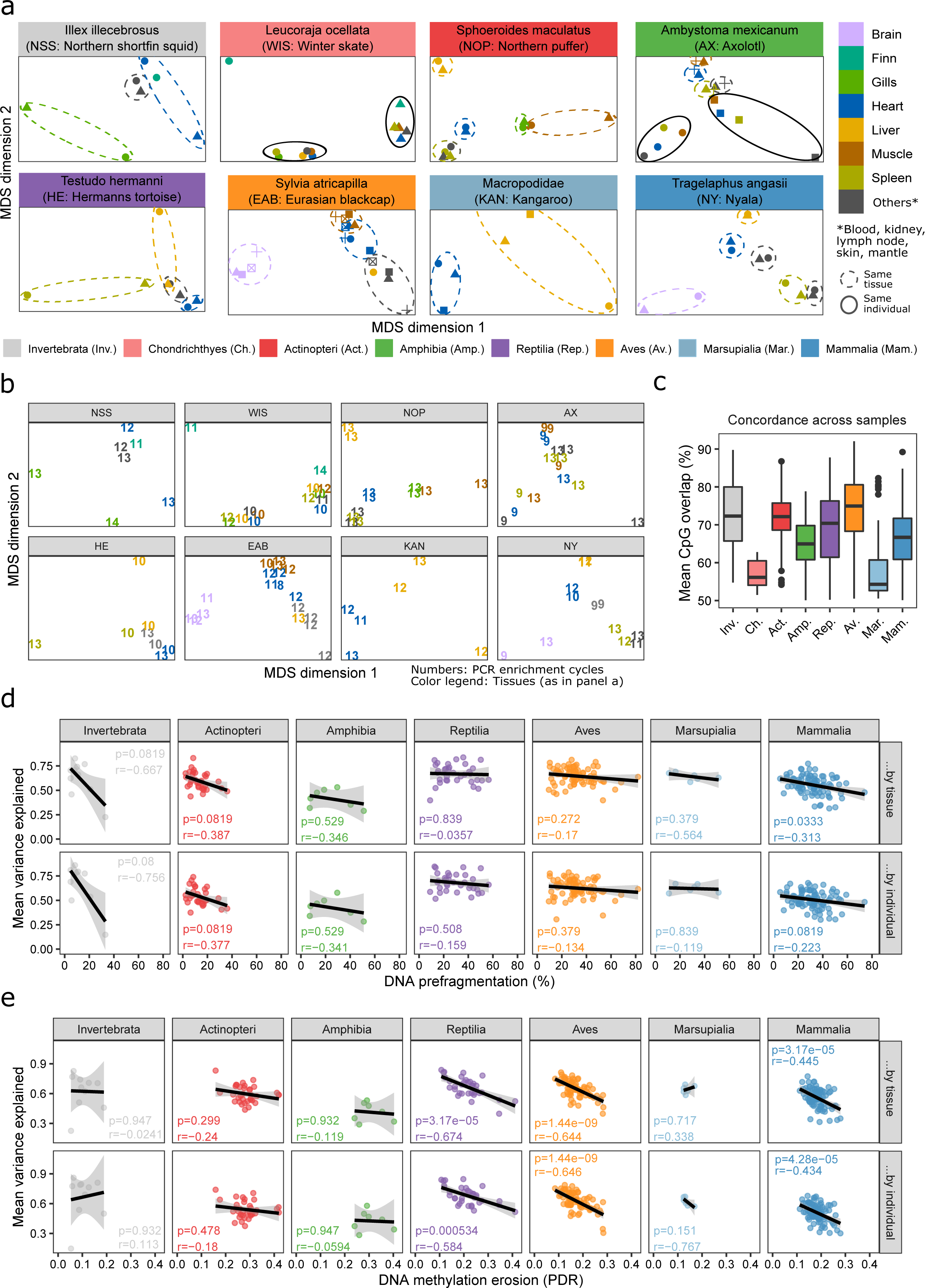
Analysis of DNA methylation profiles across tissues and individuals. (a) Multidimensional scaling (MDS) plots illustrating the similarity of DNA methylation profiles across tissues and individuals in one selected species for each taxonomic group. (b) Same as (a) but with individual data points labeled by their PCR enrichment cycles in the RRBS assay. (c) Boxplots displaying the mean overlap in covered CpGs between samples of the same species, relative to the total number of covered CpGs in each sample. This is an indicator of genetic variation in the species, in the sense that more genetically diverse samples tend to have a lower fraction of jointly covered CpGs. (d) Scatterplots relating the DNA methylation variance explained by the individual and by the tissue to the amount of DNA pre-fragmentation (as a measure of DNA quality). Each dot corresponds to one species, colored by taxonomic group. Lines indicate linear regressions for each of the taxonomic groups with 0.95 confidence intervals. Pearson’s correlation and the associated significance are indicated. (e) Scatterplots relating the DNA methylation variance explained by the individual and by the tissue to the amount of DNA methylation erosion as measured by the proportion of discordant reads (PDR). Each dot corresponds to one species, colored by taxonomic group. Lines indicate linear regressions for each of the taxonomic groups with 0.95 confidence intervals. Pearson’s correlation and the associated significance are indicted.

**Supplementary Figure 9.**
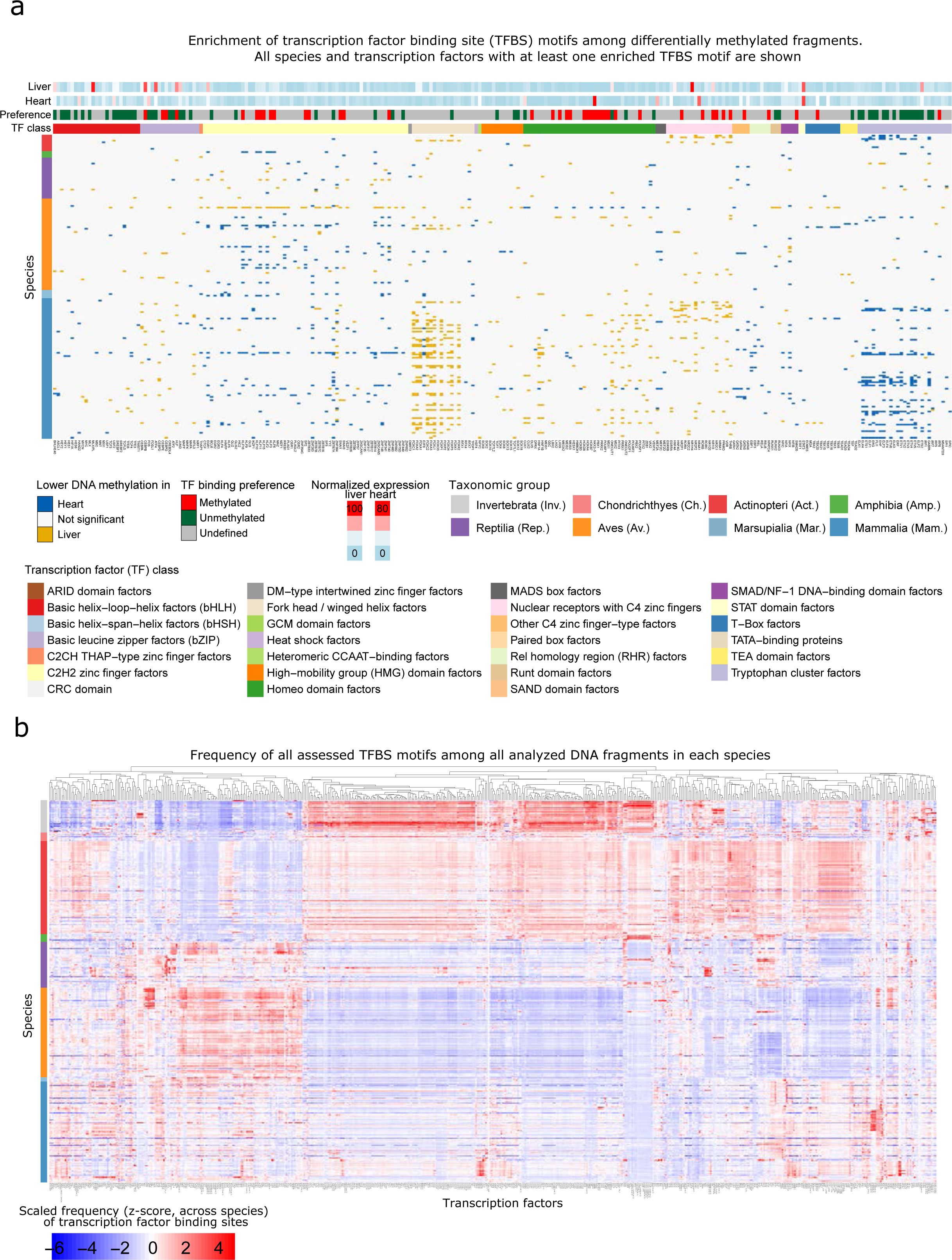
Analysis of transcription factor binding site (TFBS) motifs among differentially methylated fragments. (a) Sorted heatmap showing TFBS motif enrichments for differentially methylated fragments between heart and liver. For all species (rows) and all transcription factor (columns), the colors indicate whether the corresponding TFBS motifs were enriched in fragments that were hypomethylated in heart (blue) or in liver (yellow). The transcription factors are color-coded by binding preference and transcription factor class (top rows). (b) Clustered heatmap showing TFBS motif frequencies across all consensus reference fragments in all species (rows) and all transcription factors (columns).

**Supplementary Figure 10.**
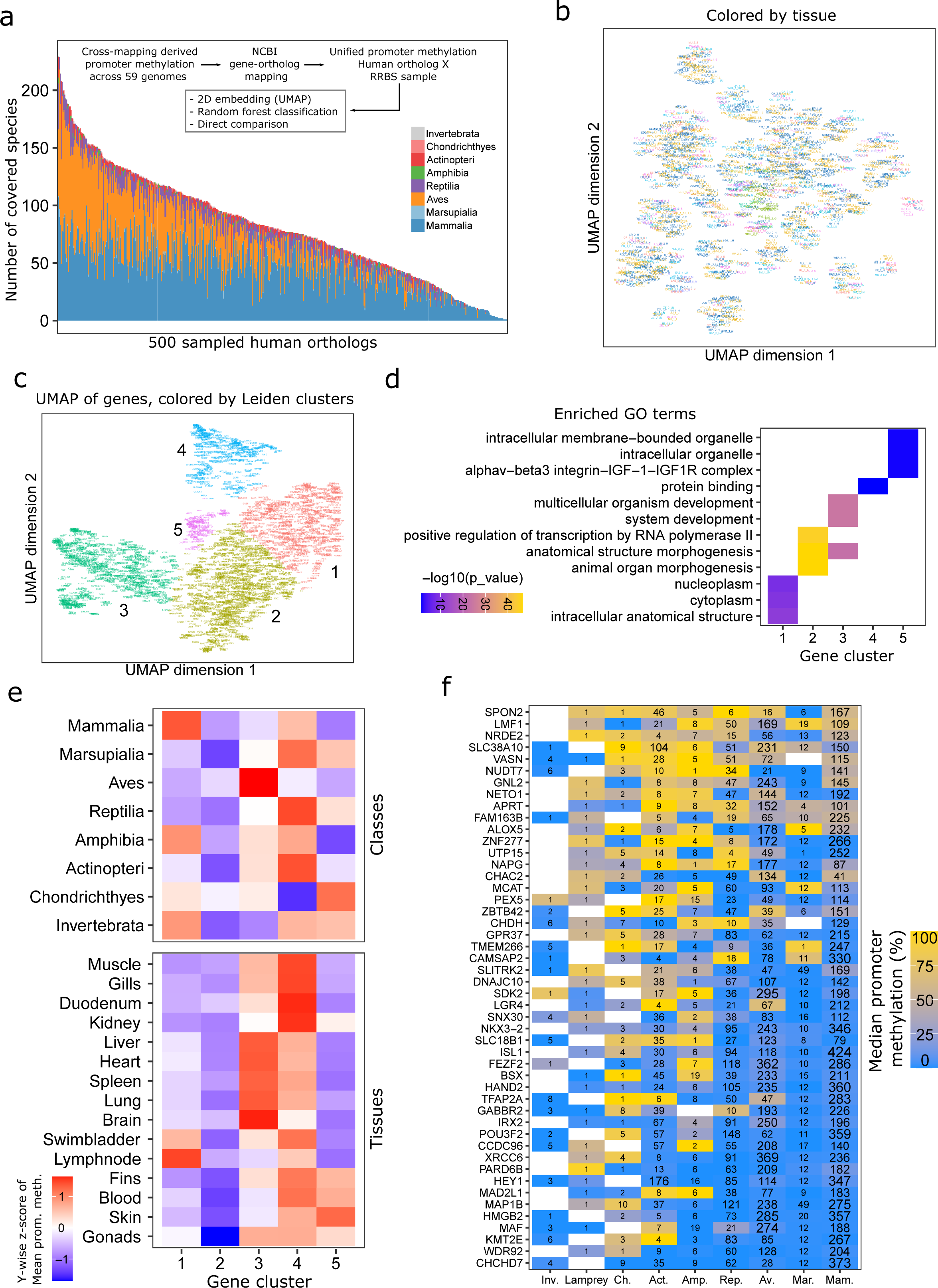
Analysis of DNA methylation at gene promoters across species in the humanortholog gene space. (a) Stacked bar plot showing the number of species per taxonomic group for specific human gene orthologs (x-axis). 500 genes were randomly samples to represent the observed spectrum. Inset: Schematic overview of the transformation of DNA methylation data into the common human-ortholog gene space and further analysis. (b) UMAP representation of DNA methylation at gene promoters based on cross-mapping of reference-free consensus reference fragments to annotated reference genomes as in Figure 5. Samples are colored by tissue, and each sample is labeled by its sample identifier (**Supplementary Table 1**). (c) UMAP representation and corresponding Leiden clustering of genes according to their promoter methylation. Genes are colored by Leiden clusters and clusters are numbered. Each gene is labeled by its name, which is searchable and readable when zooming into the PDF of the figure. (d) Heatmap showing GO term enrichments for the gene clusters defined in panel c. The top three GO terms are displayed for each gene cluster. (e) Heatmap showing scaled promoter methylation across gene clusters, taxonomic groups, and tissues, filtered at a minimum of eight samples. (f) Heatmap showing promoter methylation for genes with measurements in most taxonomic groups. Numbers correspond to the number of samples across which median promoter methylation levels were calculated.

## Supplementary Tables

Supplementary Table 1. Sample annotations and DNA methylation profiling statistics for all tissue samples

Supplementary Table 2. Overview and annotation of the animal species included in this study

Supplementary Table 3. Overview and annotation of the individual animals included in this study

Supplementary Table 4. Sample annotations and sequencing statistics for the unconverted libraries

